# Mapping satellite glial cell heterogeneity reveals distinct spatial organization and signifies functional diversity in the dorsal root ganglion

**DOI:** 10.1101/2025.05.19.654813

**Authors:** Ole A. Ahlgreen, Mads W. Hansen, Jonas Baake, Thomas E. Hybel, Rachele Rossi, Xin Lai, Ishwarya Sankaranarayanan, Johanne Pold, Lin Lin, Line S. Reinert, Søren S. Paludan, Theodore J. Price, Lone T. Pallesen, Christian B. Vægter

**Author notes:** Correspondence: Christian Bjerggaard Vægter.

## Abstract

Satellite glial cells (SGCs) envelop the somata, axon hillock, and initial axon segment of sensory neurons in the dorsal root ganglia (DRG), playing a critical role in regulating the neuronal microenvironment. While DRG neurons have been extensively studied and classified based on size, molecular markers, and functional characteristics, very little is still known about SGC heterogeneity and its potential implications on sensory processing in the DRG. Single cell transcriptional analyses have proposed the existence of SGC subtypes, yet *in situ* validation, spatial distribution, and potential functional implications of such subtypes are still largely unexplored. Here, we present the first comprehensive *in situ* characterization of SGC heterogeneity within the mouse DRG. By integrating single-cell RNA sequencing with immunohistochemistry, *in situ* hybridization, and advanced imaging techniques, distinct SGC subclusters were identified, validated, and spatially mapped within their native anatomical context. We visually identify four distinct subpopulations: 1) a predominant population of perisomatic SGC sheaths defined by the expression of marker proteins traditionally used to characterize the entire SGC population, including FABP7, KIR4.1, GS, and CX43. 2) OCT6+ SGCs occasionally being found in mosaic perisomatic sheaths, and consistently associated with axonal glomeruli, primarily ensheathing initial segment axon. 3) SCN7A+ SGCs, exhibiting no/low expression of traditional SGC markers and forming specialized homogenous sheaths around non-peptidergic neuron subtypes, implicating their potential role in pruritic (itch-related) conditions. 4) Interferon response gene-expressing SGCs, responding to Herpes Simplex Virus infection, suggesting potential involvement in antiviral protection. Finally, we investigate human DRG and find an inner perisomatic SGC layer surrounded by an outer SGC layer, with traditional and novel markers distinctively distributed between the two layers. Our results provide novel insight into SGC heterogeneity in the DRG and suggests distinct functional properties for such subtypes of relevance for the neuronal microenvironment.

## Introduction

Satellite glial cells (SGCs) are distinguished by their unique morphology, forming a thin sheath around neuronal bodies and initial segment of axons in peripheral ganglia^1,2,3,4^. This intimate contact allows a tight bidirectional communication between the glial and neuronal cells, providing SGCs with critical roles in maintaining neuronal homeostasis and regulating excitability. For the sensory neurons in the dorsal root ganglia (DRG), pathological conditions such as metabolic disease, chemotherapeutic treatment, or nerve injury can result in neuropathic pain characterized by neuronal hyperexcitability. It has been well described how such conditions also involve SGC reactivity (“gliosis”) involved in maintaining the neuropathic pain phenotype^5^ and these cells may therefore be suitable targets for pain therapy.

SGCs are characterized by specific marker proteins that have been extensively used as universal SGC markers in immunohistochemistry (IHC)^5,6,7,8,9,10,11^, hereafter termed “classical SGC markers”. These include FABP7 (Fatty Acid binding protein 7, gene *Fabp7*), GS (Glutamine synthetase, gene *Glul*), CX43 (Connexin 43, gene *Gja1*), and KIR4.1 (ATP-sensitive inward rectifier potassium channel 10, gene *Kcnj10*). Interestingly, evidence of SGC heterogeneity has emerged in recent years^12,13,14,15,16,17,18,19^, raising an important question whether such subtypes may co-existing in mosaic pattern sheaths, or if subtypes rather define homogeneous but functionally distinct sheaths. Furthermore, it is intriguing to speculate on the relationship between SGC and neuronal subtypes. Historically, murine DRG neurons have been classified as large and small diameter neurons. The large diameter proprioceptive neurons give rise to thickly myelinated afferents (Aβ) expressing neurofilament heavy chain (NFH). In contrast, the small diameter nociceptive neurons give rise to unmyelinated (C-fibers) or thinly myelinated axons (Aδ-fibers). The small diameter neurons are further subclassified into three main groups: Peptidergic, expressing calcitonin gene-related peptide (CGRP) and substance P (SP); Non-peptidergic (NP), binding isolectin B4 (IB4); and neurons expressing tyrosine hydroxylase (TH). Later work employing unbiased transcriptional profiling has resulted in more detailed subcategorization correlating with functional properties including a subset of NP that does not bind IB4^20,21^ similar advances have been achieved in the categorization of human DRG neurons, with some subsets of neurons that are likely unique in humans^22,23^

While a correlation between neuronal and SGC subtypes would add an important layer of knowledge to our understanding of sensory processing in the DRG, current evidence of SGC heterogeneity is limited to transcriptional studies. Here we advance the characterization of SGC heterogeneity by validating and correlating classical SGC markers with novel SGC markers and neuronal markers *in situ*. We identify and validate different subtypes of SGCs; 1) a widespread subtype expressing classical SGC markers and forming neuronal sheaths (hereafter termed “classical SGCs”); 2) a subtype expressing OCT6 alongside classical SGC markers, found primarily at the axonal glomerulus (“axonal SGCs”) but occasionally existing in mosaics with classical SGCs within perisomatic SGC sheaths; 3) a subtype expressing *SCN7A* but largely devoid of classical SGC markers, essentially limited to surrounding NP neurons; and 4) a SGC subset expressing *Ifit3* in addition to classical SGC markers, possibly involved in defense against viral infections targeting the DRG^24^.

## Results

### Transcriptional profiling reveals SGC heterogeneity

In this study, we analyzed single cell RNA sequencing (scRNA-seq) data from dissociated mouse DRG to identify the overall cell clustering based on the expression of known cell type mRNA markers (**Figure 1A**), as previously described^25^. Based on differentially expressed genes, the SGC cluster was divided into five subclusters of unequal proportions, ranging from the largest (Cluster 0) to the smallest (Cluster 4) (**Figure 1B, Supplementary Figure S1A**). To validate our subclustering and the identified subcluster markers, we compared our data with published studies employing scRNA-seq on sensory and autonomic ganglia^12,14,15,16,19^. Comparisons showed that the transcriptional profiles of especially our Cluster 1, Cluster 3 and Cluster 4 were highly reproducible among studies with different samples and techniques, supporting the biological relevance of the subclustering. (**Supplementary Figures S2 and S3**). We focused our validation efforts on visualization of markers that are consistently enriched in clusters in both our study and the referenced studies (**Table 1**).

**Figure 1:**
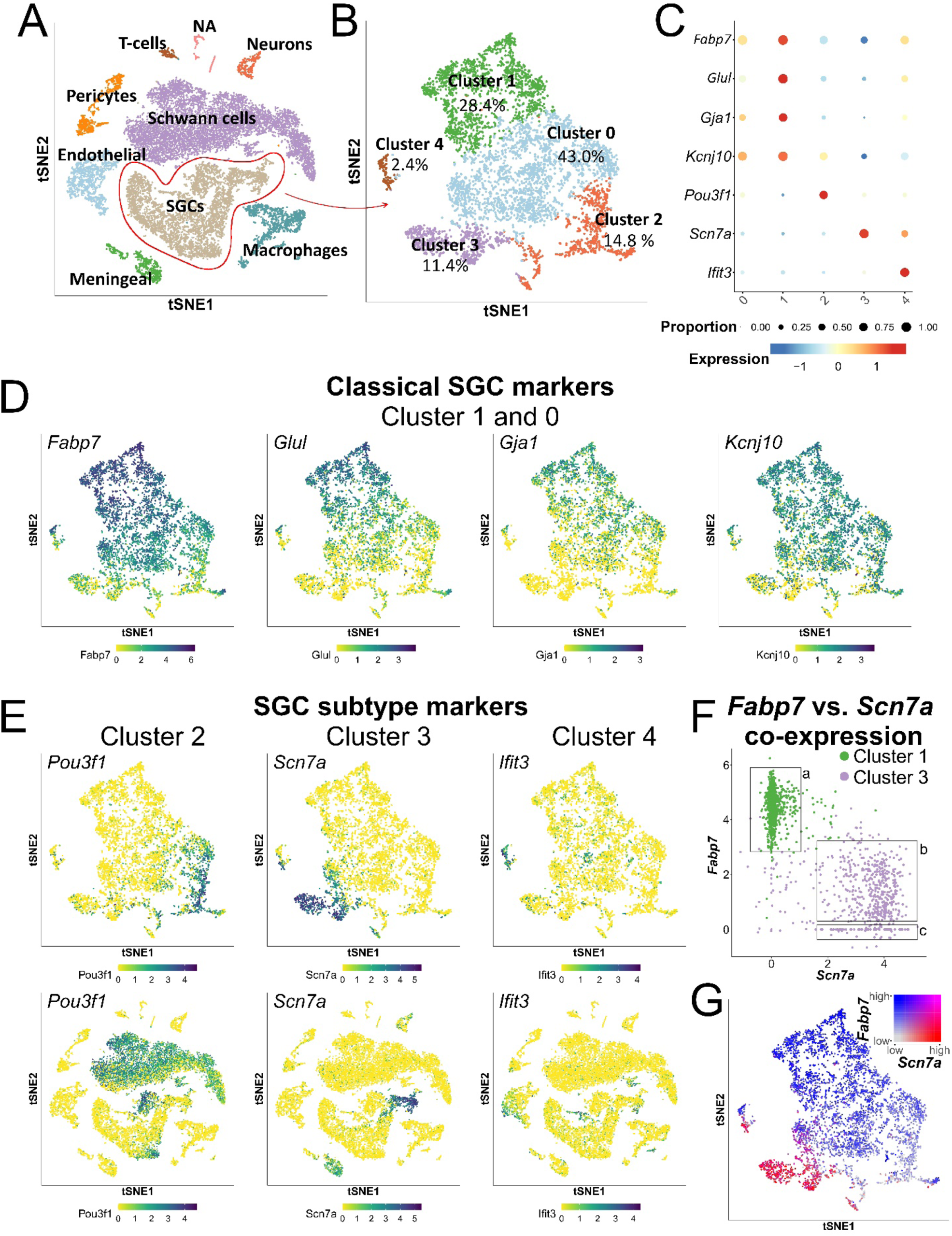
Differential expression of marker genes among SGC subclusters. **A**: Cell types within the dissociated DRG samples were annotated based on cell type marker expression. **B**: SGCs were separated into five subclusters, Clusters 0-4. **C**: Differential relative expression of marker genes among the five SGC subclusters. Cluster 0 was not meaningfully separable from the other subclusters. **D**: Cluster 1 was enriched for the classical SGC markers (*Fabp7*, *Glul*, *Gja1* and *Kcnj10*). **E**: Cluster 2 was enriched for *Pou3f1*; Cluster 3 was enriched for *Scn7a,* and Cluster 4 was enriched for interferon response genes (among these, *Ifit3*). **F**: Plotting absolute *Fabp7* and *Scn7a* expression levels among Cluster 1 and Cluster 3 cells reveals two populations among Cluster 3 cells each representing ∼50%; *Scn7a*^high^ *Fabp7*^low^ (box b) and *Scn7a*^high^ *Fabp7*^neg^ (box c). **G**: Co-expression plot shows disparate expression of *Fabp7* and *Scn7a* among SGCs.

**Table 1:** Selected enriched markers from each of the clusters of the present study are also enriched in specific clusters of other SGC heterogeneity datasets.

| **Marker** | **Ahlgreen**<br>**et al.**<br>**(present study)** | **Avraham**<br>**et al. [13]** | **Mapps**<br>**et al. [15]** | **Tonello**<br>**et al. [18]** | **Weperen**<br>**et al. [11]** |
| --- | --- | --- | --- | --- | --- |
| ***Fabp7*** | C1 | C3 | Sens, Symp | SGCs I | C2, C3 (Mature) |
| ***Glul*** | C1 | C3 | Sens | SGCs I | C1 (Immature) |
| ***Gja1*** | C1 | C3 | Sens | SGCs I | C1 (immature) |
| ***Kcnj10*** | C1 | C3 | - | SGCs I | - |
| ***Pou3f1*** | C2 | C1 | Sens, IEG | - | C4 (Aged) |
| ***Scn7a*** | C3, C4 | C4 | GR, IEG | SGCs II | C4, C5 (Aged) |
| ***Ifit3*** | C4 | C4 | Immune | - | C6 (Immune) |

Cluster 1, comprising 28.4% of all annotated SGCs, exhibits enrichment for classical SGC markers (**Figure 1C-D**) as well as genes involved in the cholesterol and fatty acid biosynthetic pathways (**Supplementary Figure S1B**) previously reported to be regulated in SGCs following peripheral nerve injury^26,27^. Cluster 2, representing 14.8% of SGCs, shows enrichment for *Pou3f1* (encoding the transcription factor OCT6), which is also strongly expressed in annotated Schwann cells (**Figure 1C and 1E**). Cluster 3, which comprises 11.4% of SGCs, is specifically enriched for *Scn7a* (encoding the sodium channel SCN7A), while Cluster 4, representing 2.4% of SGCs, exhibit enrichment of interferon response genes, with *Ifit3* encoding the interferon-induced gene IFIT3 serving as a strong marker (**Figure 1C and 1E**).

While Clusters 1-4 can be meaningfully separated based on individual marker enrichment, Cluster 0, constituting 43.0% of SGCs, cannot be readily distinguished from the other clusters using neither positive nor negative marker enrichment (**Supplementary Figure S1**). However, the expression of classical SGC markers in Cluster 0 cells (**Figure 1C-D**) albeit reduced compared to Cluster 1, aligns with Cluster 0 being assigned as part of the classical SGCs representing the vast majority of all perisomatic SGCs.

Interestingly, Cluster 3 appears to be substantially devoid of the classical SGC markers (**Figure 1C-D**). Indeed, *Fabp7* is not expressed by 49.6% of Cluster 3 cells (**Figure 1F, box c**). In the remaining 50.4% Cluster 3 cells*, Fabp7* is expressed (**Figure 1F, box b**) but at notably lower expression levels compared to Cluster 1 cells (**Figure 1F, box a**). This disparate relationship between *Fabp7* and *Scn7a* is further evident in a co-expression plot of *Scn7a* and *Fabp7* (**Figure 1G**), suggesting that Cluster 3 indeed constitutes a novel subtype of SGCs distinct from the classical SGCs.

### Spatial distribution of SGC subtypes in the DRG

The identification of SGC heterogenicity raises the possibility of distinct homogeneous sheaths composed of a single SGC subtype, as well as heterogeneous (mosaic) sheaths composed of two or more SGC subtypes. To better understand the diversity of SGC subtypes on a structural level, we aimed to visualize the novel SGC subtypes based on enriched marker genes within their identified clusters and investigate their spatial distribution in the DRG.

Cluster 2 is characterized by a distinct expression of *Pou3f1* (**Figure 1C**), encoding the transcription factor OCT6. We identified examples of OCT6+ nuclei within FABP7+ sheaths surrounding the neuronal soma (**Figure 2A**). Three-dimensional reconstruction demonstrated how such OCT6+ nuclei are completely ensheathed by FABP7+ cytoplasm (**Figure 2B**), excluding their identity as Schwann cells (FABP7-). While perisomatic OCT6+ SGCs are relatively rare events, they were consistent observed across biological replicates and suggest the presence of SGC mosaic sheaths.

**Figure 2:**
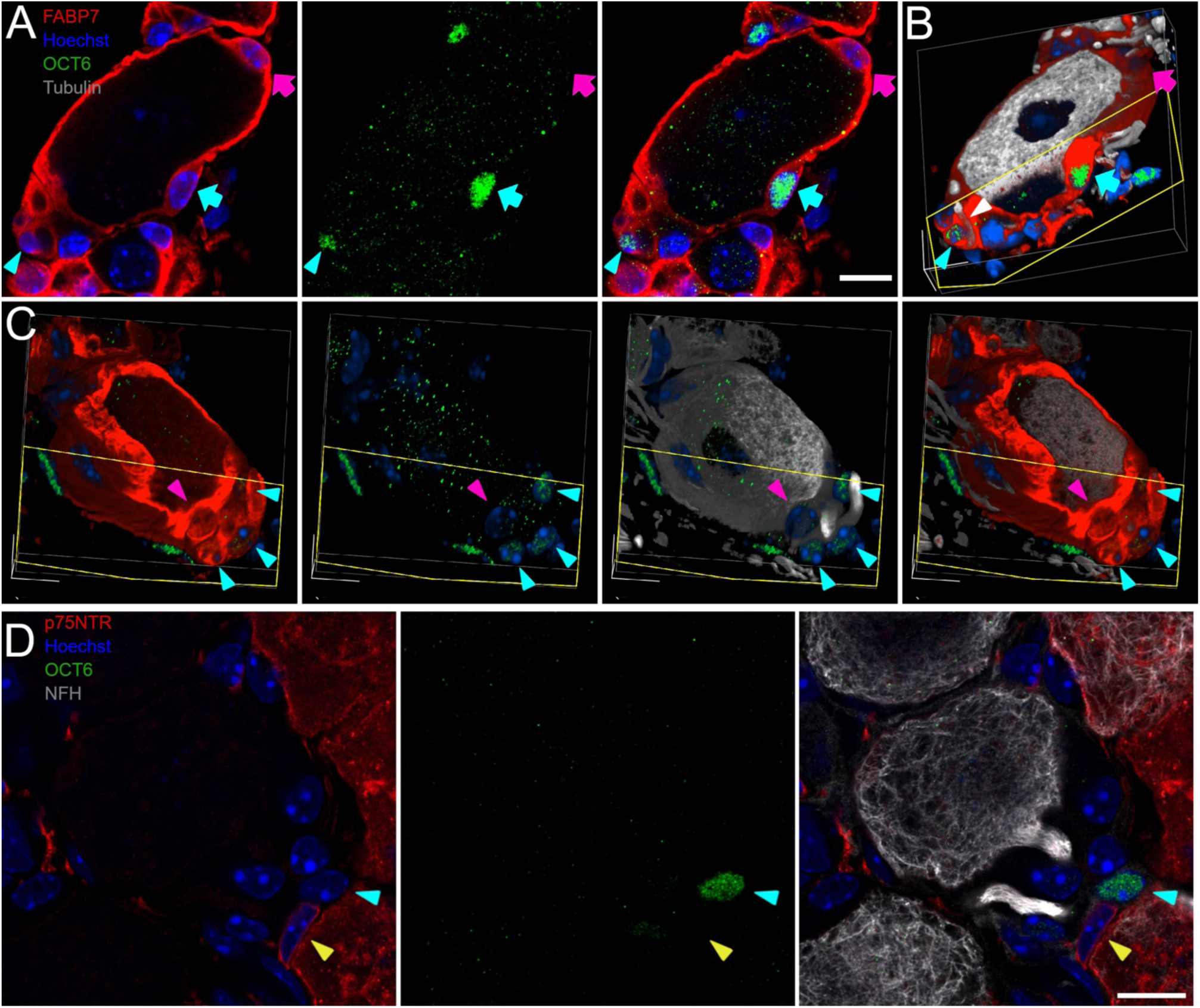
Visual representation of SGC Cluster 2 demonstrated by OCT6 expression *in situ* in lumbar DRGs. **A**: An OCT6+ nucleus (green; cyan arrow) within a FABP7+ SGC sheath (red). The same SGC sheath contains an OCT6-nucleus (magenta arrow). The axonal glomerulus of the neuron (β-III-tubulin, gray; white arrowhead) is completely ensheathed by FABP7. The FABP7 ensheathed glomerulus also contains an OCT6+ nucleus (cyan arrowhead). **B**: 3D reconstruction of a confocal z-stack of (A). The OCT6+ nucleus is completely enveloped by FABP7+ cytoplasm as shown by cross sectioning the 3D reconstruction. The initial segment axon (β-III-tubulin, gray; white arrowhead) is completely ensheathed by FABP7. **C**: 3D reconstruction of a different neuron. The FABP7 ensheathed glomerulus region contains several OCT6+ nuclei (green; cyan arrowhead). **D**: An OCT6+ nucleus (green; cyan arrowhead) in the axonal glomerulus of a large dimeter neuron (NFH, gray) is not ensheathed by p75NTR+ cytoplasm (red). In contrast, a p75NTR+ ensheathed nucleus is OCT6- (yellow arrowhead). Scalebars: 10 µm.

More prominent than the sporadic mosaic OCT6+ SGCs, we consistently observed OCT6+ nuclei completely embedded in the FABP7+ ensheathed glomerular regions containing the initial axon segments (**Figure 2**). This localization is similar to that of a previously described p75NTR++ glia cell type, found at the transitioning zone between SGCs and myelinating Schwann cells in the rat DRG ^3^. We therefore proceeded to clarify whether our OCT6+ axonal SGC are identical to the reported P75NTR++ and KIR4.1-glia. Thorough investigation of the glomerulus regions throughout the DRG revealed that OCT6+ nuclei are embedded in FABP7+ cytoplasm (**Figure 2A-C**) while p75NTR and OCT6 are positive in complementary cells only (**Figure 2D**). These findings therefore confirm that the OCT6+ axonal SGCs constitute a distinct cell population different from the described p75NTR++ glia.

Cluster 3 is characterized by distinct expression of *Scn7a* (**Figure 1C**) encoding the putative voltage gated sodium channel SCN7A. Notably, we consistently observed distinct SCN7A+ SGC sheaths surrounding a subset of neurons (**Figure 3**), with the sheaths having very low to no FABP7 co-expression in accordance with transcriptional data (**Figure 1F**). Similar SCN7A+ sheaths were also observed in mouse trigeminal and stellate ganglia, as well as rat DRG (**Supplementary Figure S4A-C**). The expression of SCN7A in non-myelinating Schwann cells prompted a deeper analysis of marker genes from our and published datasets (**Supplementary Figure S5**), which, in combination with the distinct ensheathing of neuronal somata, supports the existence cluster 3 as a defined SGC subtype. SCN7A+ SGC sheaths are consistently thinner relative to FABP7+ sheaths and can be found throughout the DRG, with a predominance towards being adjacent to the central axonal area (**Figure 3A**). Furthermore, we did not observe any mosaic sheaths presenting both SCN7A+ and SCN7A-SGCs.

**Figure 3:**
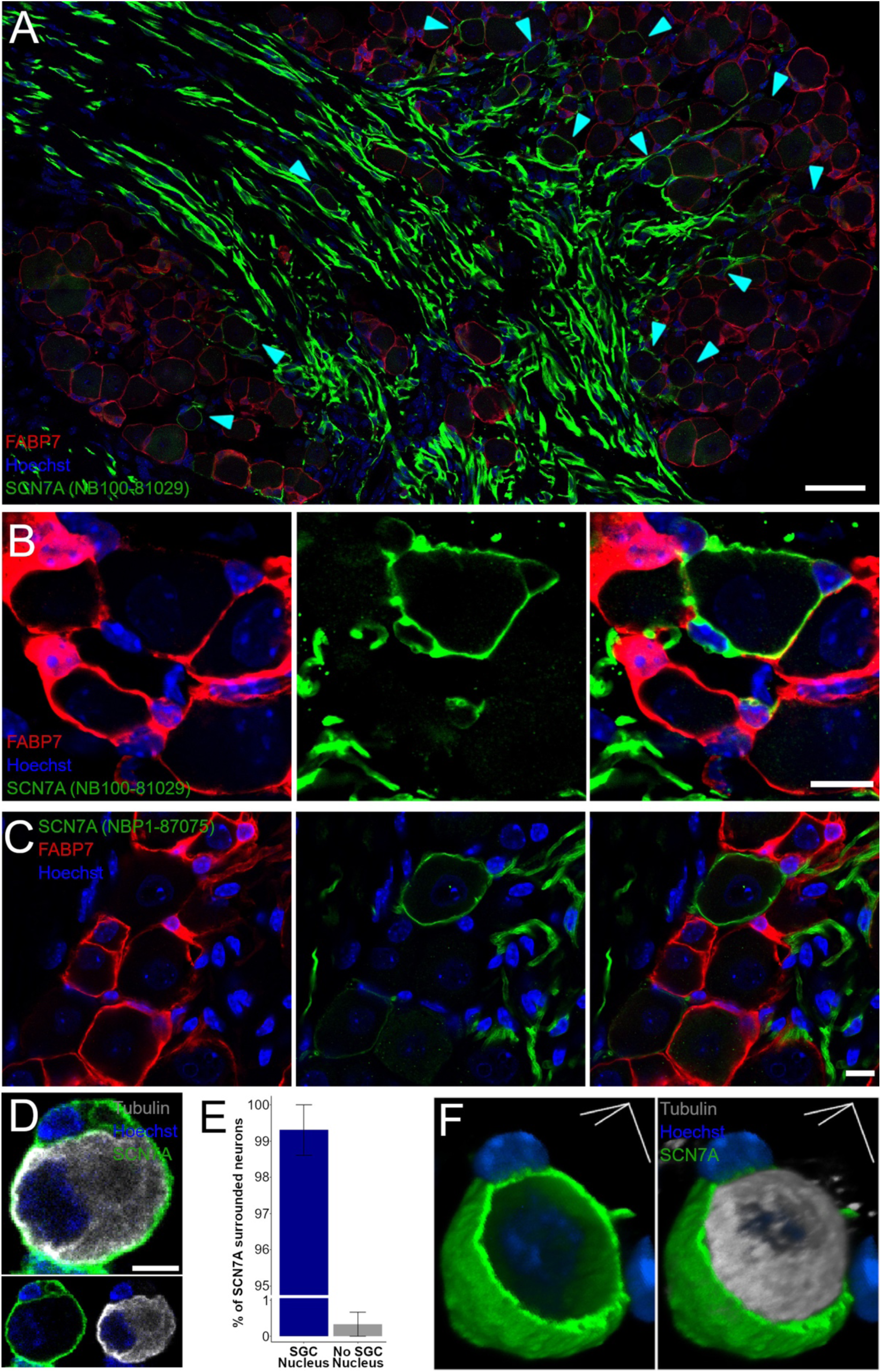
Visual representation of SGC Cluster 3 demonstrated by SCN7A expression *in situ* in lumbar DRGs. **A**: DRG section stained for FABP7 (red) showing uniform distribution of classical SGCs and SCN7A (green, cyan arrowheads) showing sparse distribution of SCN7A+ SGCs across the DRG sections with a preference toward the central axonal area. A subset of axons shows strong SCN7A reactivity. **B**: Illustration of an SCN7A+ SGC sheath with 3 ensheathed nuclei adjacent to FABP7+ SGC sheaths that are SCN7A-. **C**: Illustration of an SCN7A+ SGC sheath visualized using a different primary antibody targeting SCN7A. **D**: Example of a neuron soma (β-III-tubulin, gray) surrounded by SCN7A+ signal (green) embedding a small nucleus (blue) in a 24 hour dissociated DRG culture**. E**: Quantification of the occurrence of SCN7A+ embedded small nuclei in SCN7A+ surrounded neurons in 24 hours dissociated DRG cultures. Error bar: S.E.M. **F**: 3D reconstructed DRG neuron with associated SCN7A+ sheath in 24 hours dissociated DRG culture. Scalebars: A: 100 µm; B,C: 10 µm; D,F: 5 µm.

Given the slender appearance and proximity to the neuronal somata perimeter, we sought to exclude the possibility that the SCN7A signal is localized in the neuronal membrane rather than in the surrounding SGCs. To this end, we identified numerous examples of SCN7A+ ensheathing small perisomatic nuclei *in situ*, verifying SGC expression by two different antibodies targeting the SCN7A protein (**Figure 3B-C**). In addition, analysis of dissociated DRG cultures verified that >99% of SCN7A+ neurons were associated with SGCs (assessed as small perisomatic nuclei) (**Figure 3D-E**). 3D reconstructions of SCN7A+ SGC ensheathed neuronal somata of the dissociated DRG culture further revealed a shell-like appearance of the SCN7A positive signal of non-uniform thickness, demonstrating an outer ensheathing layer rather than a signal integrated in the neuronal membrane (**Figure 3F**). Collectively, our data demonstrate that a subset of DRG neurons are completely covered by SCN7A+ SGC sheaths, supporting the hypothesis that SGC sheaths may be functionally heterogeneous.

Cluster 4 is characterized by a distinct expression of *Ifit3* (**Figure 1C**) encoding the interferon-induced protein IFIT3. While we were unsuccessful in detecting several candidate cluster 4 marker proteins using IHC, we utilized RNAscope to visualize *Ifit3* transcripts and identified rare, but consistent, events of *Ifit3*+ perisomatic cells within the FABP7+ SGC population of naïve DRG (**Figure 4A**). Given that the interferon response is central to the innate immune system’s defense against viral infections^24^, we tested whether Herpes Simplex Virus type 2 (HSV-2) infection could increase *Ifit3* expression in cluster 4 SGCs. Examining DRGs from mice at an early stage of HSV-2 infection revealed expansive *Ifit3* expression, correlating with a cluster of HSV-2 infected neurons and SGCs (**Figure 4B-C**) but not in other areas devoid of HSV-2 infected cells.

**Figure 4:**
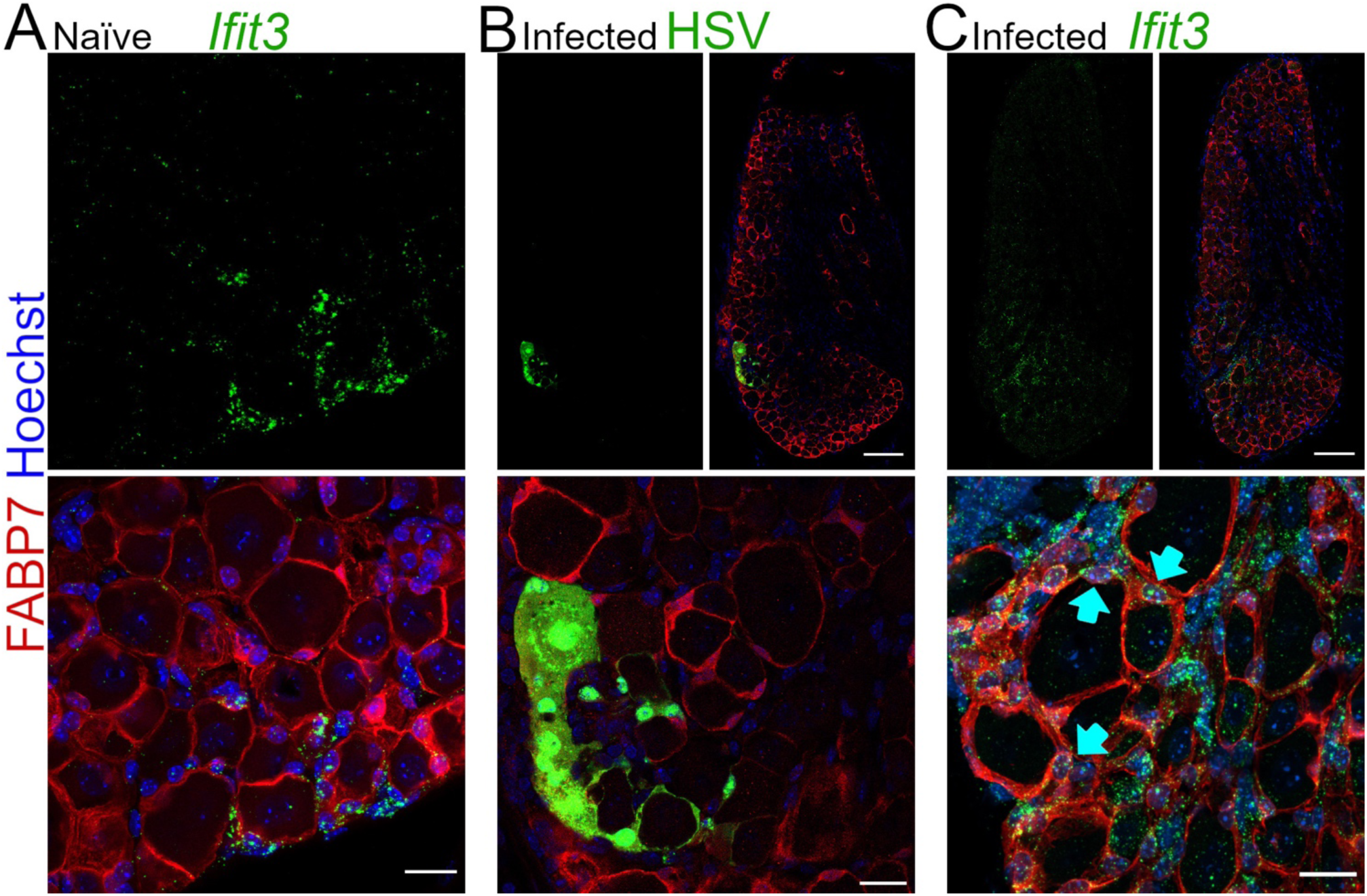
Visual representation of SGC Cluster 4 demonstrated by *Ifit3* mRNA expression *in situ* in lumbar DRGs. **A**: Occurrence of *Ifit3*+ SGCs in naïve lumbar DRG section. **B,C**: Consecutive sections of L5 DRG from HSV infected mouse. B) HSV infected cell cluster incl. neurons and SGCs within a limited area. C) Increased *Ifit3* mRNA expression in the vicinity of the HSV cell cluster from (B). *Ifit3* expressing SGCs are marked by cyan arrows. Scalebars: 20 µm in A and B,C close-ups; 100 µm in B,C overviews).

Together, our observations demonstrate the spatial distribution of SGC heterogeneity of at least four subclusters: The widespread “Classical SGCs” (here defined as SGCs expressing classical markers and forming sheaths around neuronal somata), constituting Cluster 0 and 1; Cluster 2 expressing OCT6, representing primarily axonal SGCs ensheathing the initial segment axon, and occasionally mosaic SGCs in classical perisomatic SGC sheaths; Cluster 3 expressing SCN7A, forming distinct sheaths around a neuronal subset; and Cluster 4 expressing *Ifit3*, possibly as an intrinsic property of SGCs in general related to protection against viral infection.

### Cluster 3 (SCN7A+) SGCs preferentially associate with small diameter neurons

Next, we sought to define whether the neurons surrounded by SCN7A+ SGC sheaths constitute one or more distinct DRG neuronal subpopulations. We pursued scalability by implementing the deep-learning tool deepflash2^28^, which allowed us to segment thousands of neuronal somata and correlate their characteristics with ensheathment of SCN7A+ SGCs (**Supplementary Figure S6**). First, we used deepflash2 to quantify neurons surrounded by SGCs positive for classical SGC markers FABP7, KIR4.1, CX43 and GS. Classical SGC markers label 90-95% of all neuronal SGC sheaths (**Figure 5A-B**) and show a high degree of co-expression illustrated by nearly complete overlap of FABP7+ and GS+ sheaths, with 96.1±1.5% of FABP7+ sheaths being GS+, and 98.2±0.6% of GS+ sheaths being FABP7+ (**Figure 5C**).

**Figure 5:**
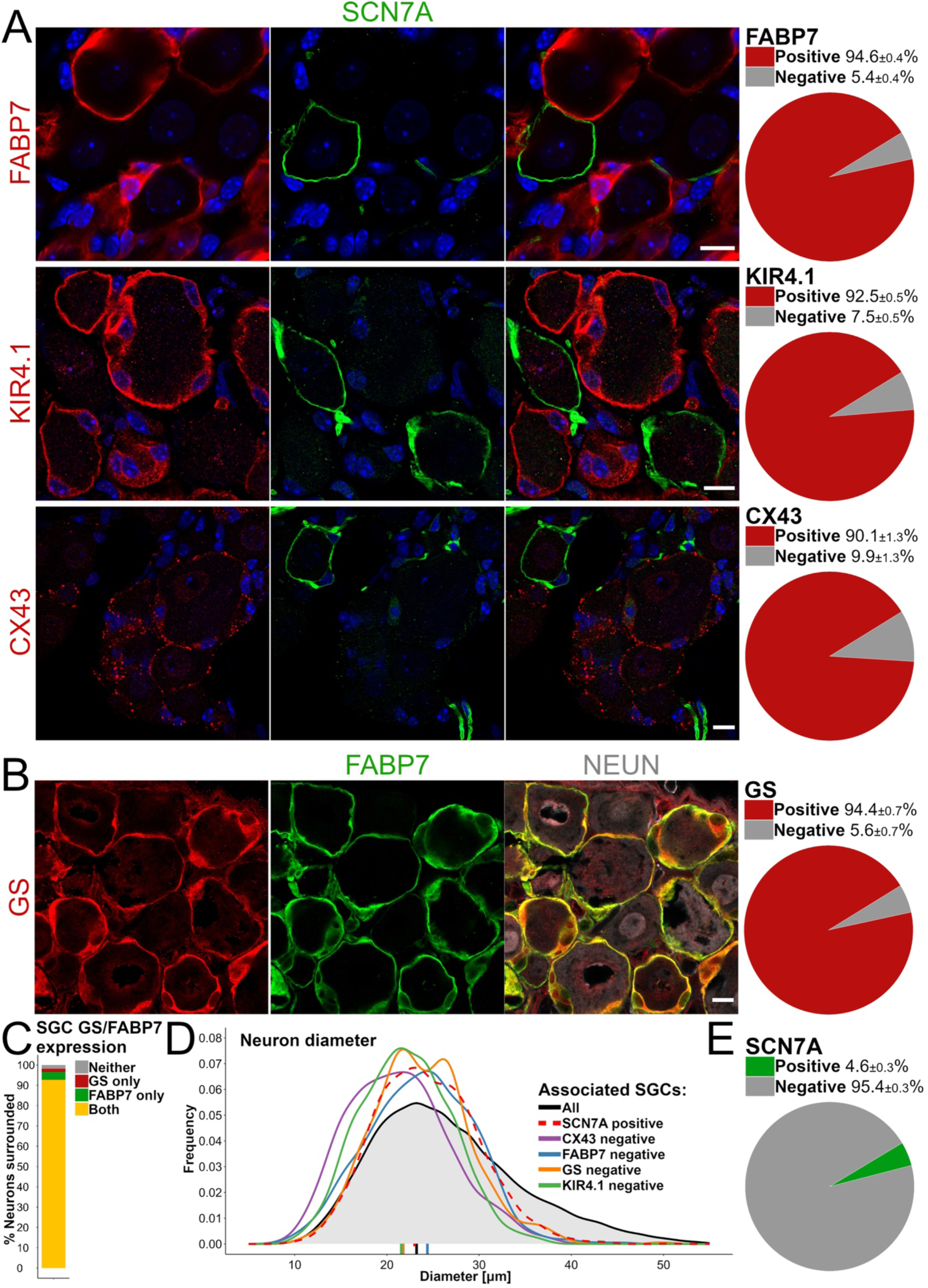
Characterizing classical SGC marker expression among SGCs and their relation to SCN7A+ SGCs. **A,B**: Representative images (left) and quantification of SGC sheaths (right, mean +/- S.E.M.) expressing classical SGC markers. Scalebars: 10 µm. **C**: Correlation of FABP7 and GS co-expression in SGC sheaths. **D**: Diameter distribution of the neurons, neurons surrounded by SGC sheaths negative for classical markers, and neurons surrounded by SCN7A+ SGCs. **E**: Quantification of the total number of neurons surrounded by SCN7A+ SGC sheaths among all neurons (NEUN+).

To determine the population size of SCN7A+ SGCs we analyzed 48,668 unique neurons (**Supplementary Table S1**) and found that 4.6% were associated with SCN7A+ SGC sheaths (**Figure 5E**). In contrast to the size distribution of classical SGC marker-ensheathed neurons which is indistinguishable from that of the total neuron population (data not shown), the size-distribution of neurons surrounded by SCN7A+ SGC sheaths is skewed towards smaller diameters and does not include the largest diameter neurons. A similar left-skewed size distribution was evident for the neurons ensheathed by SGC sheaths negative for each of the four classical SGC markers (**Figure 5D**).

### SCN7A+ SGCs preferentially ensheathe non-peptidergic neuron subclasses

The observation that SCN7A+ SGCs ensheathe small-diameter neurons suggests their association with one or more specific neuronal subtypes. Sensory neurons in the murine? DRG have been extensively studied and classified based on diameter, molecular markers, and functional characteristics. Broadly speaking, the traditional marker classification includes 1) large-diameter neurons expressing NFH, 2) small-diameter peptidergic neurons expressing CGRP and SP, 3) small-diameter NP neurons binding IB4, and 4) small-diameter TH-expressing neurons. Later works have refined these categorizations based on transcriptional profiling, revealing more than a dozen neuronal subpopulations^20,21,29^. Such subclassification has e.g. revealed an IB4-subset of NP neurons, while Plexin C1 is shown to label all NP neurons^20^. We labeled murine DRG neurons by the mentioned markers to distinguish neuronal subpopulations (**Figure 6A-B**) and applied deepflash2 to determine whether the subset of SCN7A+ ensheathed neurons represents a distinct neuronal subtype. Large-diameter NFH+ neurons represent 41.1 ± 0.8% of all DRG neurons (**Figure 6D**). However, in accordance with the smaller diameter of SCN7A+ ensheathed neurons (**Figure 5D and 6C**), NFH+ neurons are very rarely (0.7% ± 0.2%) surrounded by SCN7A+ sheaths (**Figure 6E**), and SCN7A+ sheaths rarely surround NFH+ neurons (**Figure 6F**). Similarly, TH+ neurons, as well as peptidergic neurons expressing CGRP and SP, do not demonstrate substantial association with SCN7A+ sheaths (1.4% ± 0.2% for TH, 1.1% ± 0.1% for CGRP, and 0.2% ± 0.1% for SP; **Figure 6E**). Reciprocally, SCN7A+ sheaths rarely surrounded TH+ or peptidergic neurons (**Figure 6F**).

**Figure 6:**
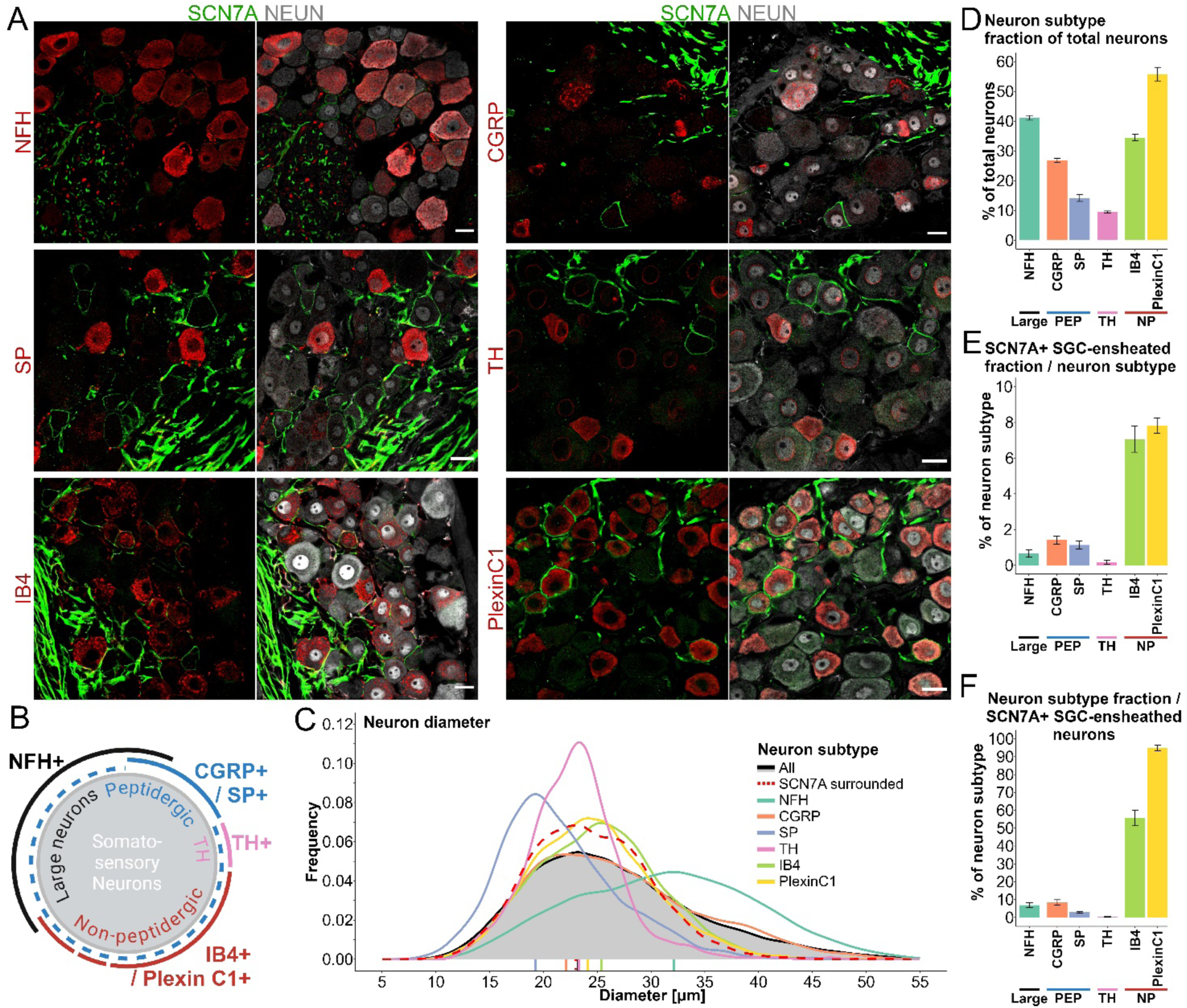
Characterizing the relationship between SCN7A+ SGC sheaths and neuronal subtypes. **A**: Representative images of SCN7A+ SGC sheaths (green) among different neuronal subtype markers (red) and the pan-neuron marker NEUN (gray). Scalebars: 20 µm. **B**: Schematic overview of neuronal subtypes within the DRG, including large-diameter neurons expressing NFH, Peptidergic neurons expressing CGRP and/or SP, TH neurons expressing TH, and non-peptidergic neurons expressing IB4 and Plexin C1. **C**: Diameter distribution of neuronal subsets based on neuronal markers, and including neurons surrounded by SCN7A+ SGC sheaths. **D**: Percentage of neuronal subtypes relative to the total number of neurons (NEUN+). **E**: Percentage of each neuronal subtype surrounded by SCN7A+ SGC sheaths. **F**: Percentage of SCN7A+ SGC sheaths surrounding specific neuronal subtypes. D-F: Mean +/- S.E.M.

In contrast, SCN7A+ sheaths appear to correlate strongly with non-peptidergic neurons, with 55.6% ± 4.4% of SCN7A+ sheaths associating with IB4+ neurons, and 94.9% ± 1.62% associating with Plexin C1+ neurons (**Figure 6F**). To further unravel the identity of NP neuronal subpopulations ensheathed by SCN7A+ SGCs, we adopted a previously described categorization; Usoskin et al. 2015^20^ define three non-peptidergic populations, named NP1, NP2 and NP3, corresponding closely to the C5+C6, C4 and C2 populations, respectively, categorized by Li et al. 2016^21^. These three classes (hereafter NP1-NP3) can be distinguished by differential expression of the markers OSMR, GFRα2 and IB4 (**Figure 7A-D**)^20,21^. We find that within the SCN7A+ SGC sheaths, 51.7% ± 2.3% are NP1 neurons, 7.8% ± 2.0% are NP2 neurons, and 35.9% ± 1.7% are NP3 neurons (**Figure 7E**). Interestingly, while SCN7A+ SGC sheaths predominantly (>95%) associate with NP neurons, mainly NP1 and NP3, only a subset of each NP1-3 subtype is surrounded by SCN7A+ SGCs (**Figure 7F**), suggesting the relevance of further functional characterization of NP1-3 neuron subtypes based on SGC sheath characteristics.

**Figure 7:**
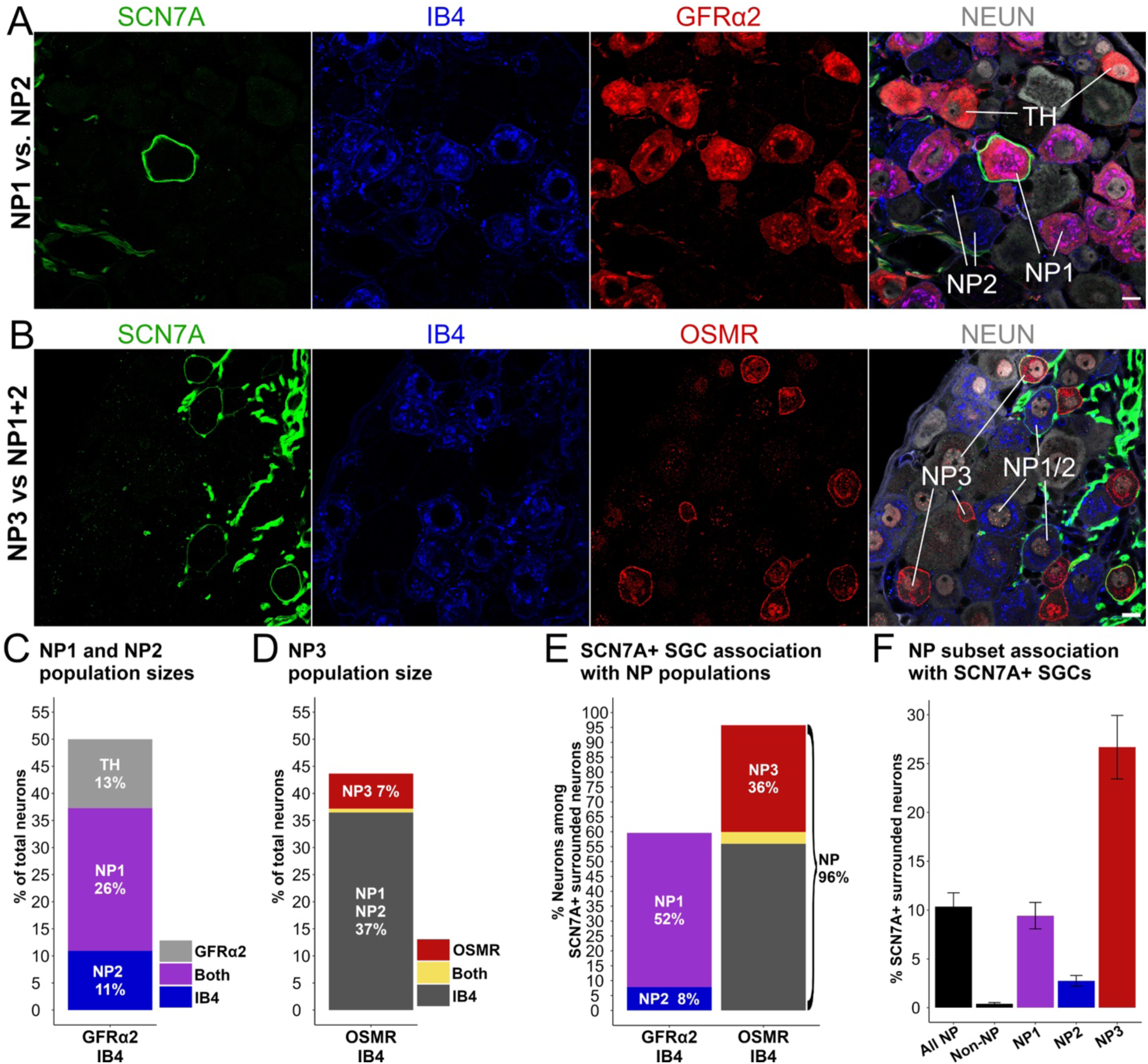
Characterizing the relationship between SCN7A+ SGC sheaths and non-peptidergic neuronal subsets. **A,B**: Representative images of DRG sections stained for SCN7A (green), IB4 (blue), GFRα2 or OSMR (red) and NEUN (gray). Neuronal subtypes were defined based on differential expression of the markers NP1: GFRα2+/IB4+; NP2: GFRα2-/IB4+; NP3: OSMR+/IB4-; TH: GFRα2+/IB4-. Scalebars: 10 µm. **C**: Quantification of NP1 and NP2 population sizes among NEUN+ DRG neurons. **D**: Quantification of NP3 population size among NEUN+ DRG neurons. **E**: Quantification of NP subsets among neurons surrounded by SCN7A+ SGC sheaths. **F**: Quantification of NP subsets being surrounded by SCN7A+ SGCs. “All NP” represent all OSMR+ and/or IB4+ neurons, whereas “non-NP” represent OSMR-/IB4- neurons. Mean +/- S.E.M.

### SGC heterogenicity in the human DRG

We extended our analysis of SGC subtypes to human DRG sections obtained from donors without indications of pain or pain-related illnesses (**Supplementary Table S2**). Using classical SGC markers applied in our mouse studies (FABP7, KIR4.1, GS and CX43), we found all neuronal somata to be surrounded by SGCs positive for all markers (**Figure 8**). Surprisingly, SCN7A also consistently ensheathed all neurons across donor tissues independent of sex and ethnicity (**Figure 8A**), representing another species difference in the DRG^23,29,30^. Furthermore, close examination of high-resolution images and 3D reconstructions demonstrated a double-layered SGC structure with an outer SCN7A+ layer of elongated or fibrous appearance surrounding an inner SGC layer (**Figure 7B**). The double layered organization appeared to correlate with soma size, as we clearly observed the structure surrounding the largest diameter neurons whereas this organization was less clear/profound surrounding smaller diameter neurons. While we regularly observed axons embedded within the outer layer or immediately adjacent to it, the SCN7A marker did not trace away from the soma along the axons, arguing that these SCN7A+ cells are not Schwann cells. The distinction between the inner and outer layers was not absolute. We observed SCN7A+ in both the inner layer and outer layer, while classical SGC markers were differently represented between the two layers; FABP7 (**Figure 7B**) and KIR4.1 (**Figure 7C**) appeared to be selective toward the inner layer, while GS was generally found in the inner layer but with some presence in the outer layer (**Figure 7D**). CX43 was clearly expressed in both the inner and outer layers, often defining two clearly separate layers (**Figure 7E**).

**Figure 8:**
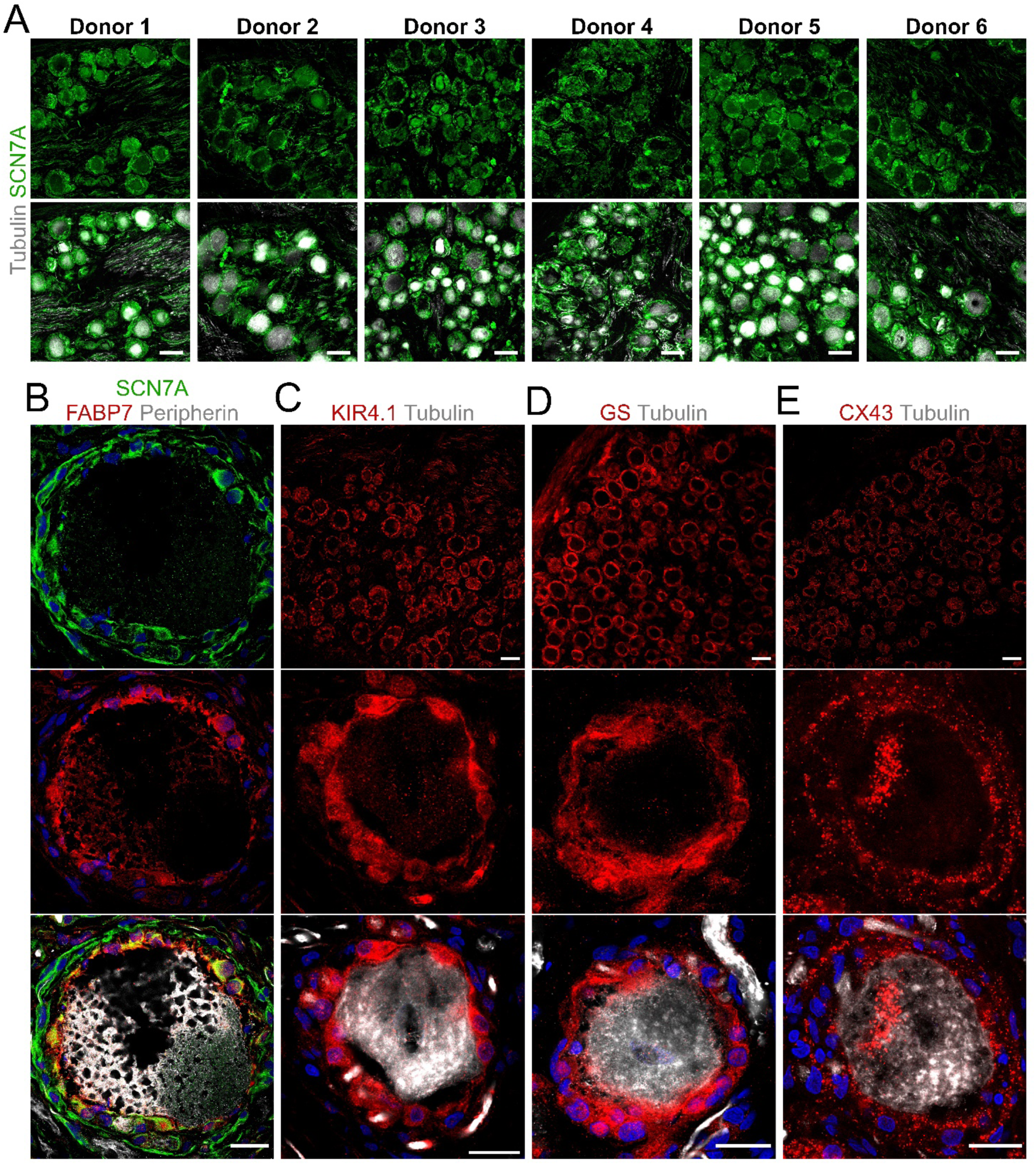
Classical SGC markers and SCN7A expression in human SGCs. **A**: Representative images of SCN7A (green) and Tubulin (gray) stains on human DRG sections from 6 donors. **B**:Close-up of a large diameter neuronal soma in a human DRG section stained for FABP7 (red), SCN7A (green) and Peripherin (gray). **C-E**: Representative images of classical SGC markers (red) including KIR4.1 (C), CX43 (D) and GS (E), all co-stained with Tubulin (gray). Scalebars: Overviews: 100 µm. Close-up single neurons: 20 µm.

These observations suggest that SGC heterogenicity exists in both murine and human DRGs. However, while single SGC sheaths are observed in mice, the largest human DRG neurons often present two concentric layers of SGCs with distinct marker profiles.

## Discussion

In this study, we mapped the heterogeneity among SGCs in the DRG using a combination of scRNA-seq, IHC and *in situ* hybridization. Our approach provides novel insights into the spatial organization of SGC subtypes, including that perisomatic SGC sheaths can exist as mosaics, being composed of more than one SGC subtype, and that SGC subtypes can form distinct homogenous sheaths, both observations pointing towards functional heterogeneity among SGC sheaths.

Using scRNA-seq we resolved the SGC population into five subclusters: Cluster 0 and 1 resemble the traditional classification of SGCs defined by various marker proteins and were therefore termed “Classical SGCs”. Cluster 2 is enriched for *Pou3f1* (encoding OCT6) among SGCs subclusters, providing a strong marker for *in situ* identification of this cluster. Cluster 3 is selectively enriched for *Scn7a*, encoding SCN7A, known as Nav2.1 in human, and Cluster 4 is enriched for interferon response genes, including *Ifit3*. We validated these clusters by comparing our data with published scRNA-seq studies^12,14,15,16,19^ and found a high degree of conservation of Cluster 1, 3 and 4 in the overall transcriptional profiles. While the Cluster 2 profile was not overall conserved among studies, the Cluster 2-enriched gene *Pou3f1* was also enriched in at least one cluster among three out of four of the compared studies. We believe that this comparative analysis highlighted the best candidate SGC markers to elucidate properties and functionalities of SGC subtypes.

Classical SGCs were defined as SGCs expressing classical SGC markers (FABP7, GS, KIR4.1 and CX43) and forming perisomatic sheaths. SGCs are believed to regulate the neuronal extracellular environment by buffering glutamate and potassium, facilitated in part by their high expression levels of GS and KIR4.1^31^. Gap junctions between SGCs are less prominent in normal conditions but become increasingly apparent in pathological conditions^32^ and are predominantly composed of CX43^33^. SGCs have also consistently been found to be enriched for metabolic pathways of lipids including cholesterol and fatty acids^10,26^, which in our data was characteristic for Cluster 1 SGCs and (albeit at a relatively lower level) in Cluster 0 SGCs. Notably, classical SGC markers, as well as enrichment for proteins involved in cholesterol biosynthesis, are well conserved in SGCs between several species including mouse, rat, human, guinea pig, and macaque^10,34^, suggesting important evolutionary conserved functions to SGCs. We show by IHC that classical SGCs constitute 90-95% of all perisomatic SGC sheaths, confirming that the classical SGC subtype represents the main contributor to the perisomatic SGCs population. Classical SGCs have therefore collectively represented SGCs in the literature, largely because SGC studies usually rely on one or more of the classical SGC markers. As a result, the classical SGC subtype is relatively well characterized. However, the remaining 5-10% of the perisomatic SGC population cannot be characterized using the classical SGC markers and presents an interesting opportunity to understand the functional implications of heterogeneity among SGCs.

Alongside the classical SGC morphology where SGCs form thin sheaths surrounding neuronal somata, axonal SGCs ensheathe the initial segment axon which forms a glomerulus located immediately adjacent to the soma^1,2,3^. Based on *Pou3f1* enrichment in Cluster 2 SGCs of the present study, as well as enrichment in at least one cluster among three out of four compared scRNA-seq studies^12,14,16^, we validated OCT6 as a marker. OCT6 is a transcription factor involved in the initiation of myelination in the developing nervous system^35^. However, in adult mice, the continued expression of OCT6 in MPZ expressing Schwann cells was found to lead to hypomyelination^36^. It is therefore possible that OCT6 is expressed in certain non-myelinating cells such as SGCs. We identified mosaic SGC sheaths around neuronal somata. These sheaths contain both OCT6+ and OCT6-SGC nuclei within the same FABP7+ sheath, and although relatively rare events, this pattern suggests the possibility of functional heterogeneity within individual SGC sheaths. The precise nature of such functionality remains speculative but may involve special interactions with surrounding structures.

While perisomatic OCT6+ SGCs cannot account for the population size of Cluster 2 cells (∼15% of the total SGC population), we also observed OCT6+ nuclei within the FABP7+ sheath surrounding most axonal glomeruli. Unlike the perisomatic SGCs, the axonal SGCs have not received much attention. One study demonstrated the presence of SOX2 in both perisomatic SGCs and axonal SGCs of large caliber (NFH+) neurons^2^. Another study identified a glial cell type located around the axon initial segment, strongly expressing p75NTR (encoded by *Ngfr*) in the rat DRG. These cells, termed p75NTR++ glia, express the pan-glia markers S100 and Sox10 and loosely cover other glial cells surrounding the initial axon segment, including axonal SGCs (p75NTR- and KIR4.1+) and myelinating Schwann cells (P75NTR-, MPZ+ and MBP+)^3^. We examined the axon initial segment glomeruli in sections stained for p75NTR and OCT6 and found that OCT6+ nuclei are not embedded in p75NTR+ cytoplasm, identifying them as axonal SGCs different from p75NTR++ glia.

A common finding among SGC heterogeneity studies is a *Scn7a*-enriched cluster (Cluster 3 in the present study). Interestingly, we visually identified a subset of SCN7A+ SGCs *in situ* forming thin homogenous sheaths surrounding approximately 5% of all neuronal somata, distributed throughout the DRG with a preference of adjacency to the central axonal area. Approximately half of these sheaths are negative for classical SGC markers, with the other half being only faintly positive for FABP7. Lacking other classical SGC markers such as KIR4.1 and CX43, the SCN7A+ SGCs might act differently regarding potassium buffering and gap junction-mediated communication observed in classical SGCs, and may not present SGC hallmarks of neuropathic settings such as KIR4.1 downregulation and increased gap junction connectivity^37,38,39^.

Our novel identification of heterogenous SGC sheaths within the DRG brings with it the potential for functional heterogeneity, with distinct SGC-neuron subtype combinations. We found a high correlation between SCN7A+ SGC sheaths and the pan-NP marker Plexin C1^20^. The individual NP subpopulations can be closely approximated by differential expression of IB4, GFRα2 and OSMR; NP1-neurons expressing IB4 and GFRα2; NP2-neurons expressing IB4 but not GFRα2; and NP3-neurons expressing OSMR but not IB4^20,21,40,41^. Together, these NP subpopulations accounted for 96% of neurons surrounded by SCN7A+ SGC sheaths, providing strong correlation of SCN7A+ SGCs specifically with NP neuron subsets. This correlation in turn suggest the relevance of SCN7A+ SGCs in relation to specific sensory functionalities, particularly pruritus (itching), since all three NP neuron subcategories are involved in pruriception^42,43,44,45,46,47,48^. Furthermore, while only a subset of each NP neuron subcluster is ensheathed by SCN7A+ SGCs, it is intriguing to speculate how specialized SGC sheaths may contribute to functional heterogeneity within clusters of sensory neuron subtypes, arising from variations in glial modulation.

Another finding in common between the compared scRNA-seq studies is the existence of a small population of SGCs expressing interferon response genes^12,16^. To validate our Cluster 4 interferon response genes-enriched SGC subpopulation, we unsuccessfully tested six different antibodies in DRG sections, with the lack of signal presumably arising from interferon protein expression below the detection threshold. We therefore focused our efforts on cluster 4 validation by *Ifit3* RNAscope, in line with transcriptional data. This revealed rare occurrences of *Ifit3*+ SGCs, as defined by colocalization with perisomatic FABP7+ SGC sheaths. Furthermore, we found that *Ifit3* was specifically induced in DRG areas corresponding to the location of HSV-infected neurons and SGCs. The correlation between HSV-2 infection and induced *Ifit3* expression was underlined by the lack of *Ifit3* expression in absence of infected cells. While the rare occurrences of *Ifit3* expressing SGCs in the naïve DRG support a specialized subtype, the inducibility may also suggest dynamic immune properties of SGCs, partaking in immune surveillance and innate immune response within the DRG. Future studies will be required to study the dynamics of SGC *Ifit3* induction *-* or other interferon response genes – at different timepoints of HSV-2 infection, as the interferon response might be limited to a certain time window or correlate more strongly with increased number of HSV infected DRG cells. This could be extended to other viruses with the capacity to infect DRG sensory neurons under experimental settings such as the Zika virus^49^ and Rabies virus^50^, swine hemagglutinating encephalomyelitis virus shown to be engulfed by SGCs when released from neuronal somata^51^, and SARS-CoV-2^52^. SGCs are often considered to form a protective barrier around the neuronal somata, but in the case of viral infections arriving to the DRG through axonal transport, it may be speculated how SGCs can constitute a barrier protecting the extracellular environment to limit the spread of infection.

Interestingly, a recent study identified the existence of microglia-like cells in the DRGs of certain species, including humans, but not in mice and rats^53^. In humans, these microglia-like cells enwrap the neuronal soma within the SGC sheath^53^. Although they are clearly a distinct cell population and not specialized SGCs (based on their transcriptional profile and the absence of classical SGC markers, such as GS), this human tri-cellular structure - comprising neuron, SGC and microglia-like cells - suggests important immune-related species differences between humans and rodents^53^. Our data suggests that in the mouse DRG, SGCs are involved in the anti-viral response, and previous studies have demonstrate that resident macrophages in mouse DRG become activated and proliferate following nerve injury^54^, even appearing to invade the SGC-neuron unit^26^. The immune functions of microglia-like cells in the human DRG and involvement of DRG resident macrophages remain to be explored but may highlight important translational considerations when using rodent model systems, for example in pain research. In fact, the vast majority of SGC studies have been conducted in rodent DRGs whereas little is still known about structural and functional characteristics of SGCs in human DRGs. We here show that in human DRGs, SCN7A is expressed in SGCs surrounding all neurons (along with classical markers), and that SCN7A+ SGCs for large-diameter neurons form a fibrous-like outer SGC layer distinct from an inner neuron-ensheathing layer of SGCs, the latter expressing classical SGC markers. While microglia-like cells were recently reported to also envelop the neuronal soma beneath the SGC sheath^53^, the presence of classical SGC markers in the inner layer does not suggest that this inner layer is composed of microglia-like cells rather than SGCs, supporting the existence of a more complex SGC organization in human DRG relative to that of mice. This layered organization, which was particularly evident for large-diameter neurons, may reflect a more complex and specialized glial architecture and functionality in humans compared to rodents. Notably, human neurons are substantially larger than mouse neurons^53,55,56^ and likely also hold higher metabolic demands. The two layers for large-diameter human neurons may be electrically and metabolically coupled, as demonstrated by a comparable layered organization of the gap junction protein CX43. While expression of SCN7A was also observed along axons (presumably in Schwann cells), the abundance was much greater around somata. Importantly, the SCN7A signal surrounding the soma did not appear to follow axons away from the soma, and the signal was present around the soma irrespective of the SGC layer having embedded axons within. Together, these observations support that the neuron-ensheathing SCN7A+ cells are indeed SGCs and not axon-associated Schwann cells.

Conservation across species generally suggests a fundamental role in sensory processing that has been maintained throughout evolution. The transferability of results from mice to humans is an advocate for the potential usefulness of further exploring the SCN7A+ SGCs, yet the differences in organization between rodents and humans also highlight the importance of considering species-specific variations when studying SGCs with a translational perspective.

In conclusion, we demonstrate for the first time the spatial distribution of murine SGC subtypes *in situ*, providing a framework to explore their roles in health and disease. We describe: “Classical SGCs” expressing classical SGC markers including FABP7, KIR4.1, GS and CX43, representing 90-95% of all perisomatic SGC sheaths; OCT6+ SGCs ensheathing the initial segment axons and occasionally involved in mosaic perisomatic SGC sheaths; SCN7A+ SGCs with low/no classical SGC marker expression, forming distinct sheaths specifically around NP neuron subsets implicating a role in pruritus; and an *Ifit3*+ SGC subset expressing interferon response genes, with implications in protection against viral infection. Our findings emphasize the relevance of understanding glial heterogeneity in relation to the neuronal microenvironments and may open for new therapeutic targets in sensory and immune-related conditions.

## Methods

### Animals

Wild type male and female C57Bl/6JRj mice were purchased from Janvier Labs (France). Mice were group housed with water and chow available ad libitum in a 12:12 hour light-dark cycle. Animals, 12-25 weeks old, were euthanized by transcardial perfusion with PBS, while deeply anesthetized with isoflurane. DRGs were accessed by cutting the vertebral column enclosing the spinal cord bilaterally from the upper cervical region to the sacrum. Specific DRG were identified by counting from costae^57^. Experiments were approved by the Danish Animal Experiments Inspectorate under the Ministry of Justice (permission no. 2021-15-0201-00967 and 2021-15-0201-01084) and carried out according to the European Council directive, and institutional and national guidelines.

### Human DRG recovery

All human tissue procurement procedures were approved by the Institutional Review Board at the University of Texas at Dallas. Human DRGs were surgically extracted using a ventral approach^58^ from organ donors within 4 hours of cross-clamp and placed immediately on artificial cerebrospinal fluid (aCSF). All tissues were recovered in the Dallas area via a collaboration with the Southwest Transplant Alliance as described in a previously published protocol^59^

### HSV infection

For vaginal HSV-2 infection, mice were pretreated by subcutaneous injection of 2 mg Depo-Provera. Eight days later, mice were anesthetized with isoflurane and inoculated intravaginally with 20 μl HSV-2 (6.7 × 104 PFUs) in PBS. Subsequently, mice were placed on their backs and maintained under anesthetics for 10 minutes. At 5 days post infection, mice were sacrificed as described above for harvesting DRG.

### Immunohistochemistry

Following PBS perfusion, mice were perfusion fixed with 4% PFA. Freshly dissected ganglia were submerged directly in 4% PFA and incubated 1.5 hour at room temperature followed by overnight cryoprotection in 30% sucrose at 4°C. Tissues were embedded in O.C.T compound and frozen solid on dry ice, sectioned at 12 µm using a Leica CM3050S Cryostat and thaw mounted onto Superfrost^TM^Plus adhesion glass slides. Slides were thawed at room temperature and rehydrated in PBST (PBS with 0.15% Triton X-100). Sections were blocked in 4% normal donkey serum in PBST for 1 hour at room temperature, incubated with primary antibodies overnight at 4°C and next day, incubated at room temperature for 4 hours with secondary antibodies and 10 min with Hoechst (H3570, Invitrogen). Coverslips were mounted with fluorescent mounting medium (Dako, S3023). For antibody and staining reagent concentrations, see **Supplementary table S1.**

### In situ hybridization

*In situ* hybridization was performed in combination with immunofluorescence in accordance with the guidelines provided in the user manual “RNAscope™ Multiplex Fluorescent Reagent v2 Assay combined with Immunofluorescence - Integrated Co-Detection Workflow (ICW)” for fixed-frozen tissue. In brief, sections were heated to 60 °C for 30 min, fixed with 4% PFA, and stepwise dehydrated in EtOH (50%, 70%, and 2x 100%) followed by incubation with H₂O₂. Target retrieval was performed prior to overnight incubation with primary antibodies diluted in 4% donkey serum at 4 °C. The following day, sections were post-primary fixed in 4% PFA and incubated with protease IV (ACD, 322331) for 30 minutes at 40°C. Probe hybridization was performed by incubating for 2 hours at 40 °C followed by sequential incubation with amplification reagents (ACD 323101, ACD 323102, and ACD 323103), HRP reagents (ACD 323106), and the TSA Vivid™ Fluorophore 520 (ACD 323271). Sections were then incubated with secondary antibodies for 2 hours and with Hoechst (H3570, Invitrogen) for 10 minutes at room temperature. Finally, coverslips were mounted using fluorescent mounting medium (Dako, S3023). For probe information and antibody and staining reagent concentrations, see **Supplementary table S1.**

### Dissociated Dorsal root ganglion cell culture

From each mouse, 40-50 DRGs were collected and conserved in ice-cold HBSS (H6648, Sigma Aldrich). Sequentially DRGs were incubated with Papain (13.3 U/ml; #LS003126, Worthington Biochemical Corporation) diluted in 3 ml HBSS with 3 µl NaHCO_3_ saturated H₂O and 1 mg of L-Cysteine for 30 minutes at 37°C, 5% CO2, and with Collagenase II (1040 U/ml; #LS004176, Worthington Biochemical Corporation) and Dispase II (5 U/ml; D4693-1G, Sigma Aldrich) diluted in 2 ml DPBS (Thermo Scientific, SH3002802) for 30 minutes at 37°C, 5% CO2. Tissues were then washed in 2 ml culture media (DMEM/F12, D8437, Sigma Aldrich (v/v) with 10% Fetal Bovine Serum, 1% Penicillin/Streptomycin) and resuspended in 1 ml culture medium supplemented with 25 ng/ml NGF and 1 μl DNase and were gently triturated and mechanically dissociated until the solution gained a homogenous aspect. Cell suspensions were then filtered through a 100 µm cell strainer and plated on coverslips coated with laminin (0.2 mg/ml, L2020, Sigma Aldrich) and 2 hours poly-L-lysine (0.05 mg/ml; 8920, Sigma Aldrich). 24 hours after plating, cells were fixed in 4% PFA for 10 minutes. Immunocytochemistry was performed as described in the immunohistochemistry section, with secondary antibody incubation of only 1 hour at room temperature.

### Single cell RNA sequencing analysis

L3 and L4 DRGs were collected in ice-cold HBSS (Gibco, 14170088) from both sides of two mice per group, totaling eight DRGs per sample across five groups. DRGs were centrifuged at 500 × g for 4 minutes at 4°C, then incubated in dissociation buffer (2.5 mg/ml collagenase, Sigma C9722, and 5 U/ml dispase II, Sigma D4693, in DPBS) at 37°C in 5% CO₂ for 30 minutes. The tissue was triturated with a P1000 pipette until homogeneous, followed by the addition of 9 ml PBS and centrifugation at 500 × g, 4°C for 8 minutes. Next, cells were incubated with 0.5 ml trypsin-EDTA (Sigma 59418C) at 37°C for 10 minutes, neutralized with HBSS containing 10% FBS (Sigma F9665), and centrifuged again. The pellet was resuspended in HBSS with DNase I (40 Kunitz units, Sigma DN25-1G) and filtered through a 40 µm strainer (VWR 734-0002). After a final centrifugation at 500 × g, 4°C for 10 minutes, cells were resuspended in PBS with 5% FBS at 1000 cells/µl.

For scRNA-seq, cell suspensions were prepared using the Chromium Single Cell 3’ GEM, Library C Gel Bead Kit v3 (10x Genomics, PN 1000075). The 10X Chromium device encapsulated individual cells into droplets with unique sequencing barcodes. Libraries were sequenced using the DNBSEQ-G400 platform, and raw reads processed with Cell Ranger v3.0.2, aligned to the mm10-3.0.0 genome (Ensemble 93). To minimize batch effects, all libraries were prepared on the same 10X chip. Count matrices were analyzed in Seurat v3 (R), filtering cells with <200 genes or mitochondrial gene expression >30%. DRG cells were annotated using marker genes, with further clustering applied to cells identified as SGCs.

### Image acquisition and processing

Images were acquired using a Zeiss LSM800 Confocal microscope. For quantifications, images were acquired as tile-scans of complete DRG sections with a Plan Apochromat 40x/1.4 Oil objective using four lasers (405 nm, 488 nm, 561 nm, 640 nm). The pin hole was adjusted for equal focal thickness of all wavelength and acquisition settings at a resolution of 70nm/pixel and additional parameters were kept constant for each stain combination. FIJI^60^ was used for all image analyses, including pre- and post-processing of images and ROIs prior to and following deepflash2 predictions. The 3DScript plugin^61^ was used for 3D rendering of confocal z stacks.

### Deepflash2 segmentation and quantification

Deepflash2^28^ was applied to create instance segmentation of neuronal somata from the NEUN channel of stained DRG sections. Initially, three annotators annotated neuronal perimeters on 7 tile scan images, which were resized to 50% alongside the corresponding NEUN channel images. The mask sets were used to establish the ground truth mask of each image using simultaneous truth and performance level estimation (STAPLE)^62^. The ground truth masks were used with the same resized images of NEUN to train a five-model prediction ensemble. Deepflash2 was applied using default parameters to predict neuron masks of 704 NEUN images originating from various multi-channel images.

For instance segmentation, deepflash2 implements the Cellpose library^63^ generating regions of interest (ROIs) compatible with ImageJ. Output ROIs were well fitted on neuronal somata and aside from occasional erroneous pixel sized ROIs, which were removed, ROIs were otherwise unaltered. ROIs were indexed, and these indices, providing the total number of neurons from each image and spatial parameters incl. area and diameter, of each individual neuron were used for relative quantification of all other channels of individual images. One annotator carried out all SGC and neuronal marker annotations semi-automatically using pixel intensities and their standard deviation using ImageJ, blinded to the neurons surrounded by SCN7A+ SGCs. The other annotator manually annotated all neurons surrounded by SCN7A+ SGCs, blinded to the SGC or neuronal marker to be correlated and including whether a particular were annotated as positive.

Quantification of the frequency of SCN7A+ SGCs were performed by combining individual datasets derived from images of sections spaced at least 60 μm apart, since diameter analysis of all segmented neurons showed that only 27 out of 154.821 (0.017%) had a diameter greater than 60 μm. This dataset includes 48,668 unique neurons across 214 non-overlapping DRG sections, from a total of 14 mice. Marker information from both annotators was added to the ROI indices and the data exported for analysis. For sample information on individual datasets produced, see **Supplementary table S2**.

### Data analysis and statistics

Data analyses, statistics and plotting was performed using RStudio version 2023.3.0.386^64^ with base-r functions and the Tidyverse^65^ package. The Wilcoxon Rank Sum test was used to compare clusters and conditions within the scRNAseq dataset. An unpaired two-tailed t test with equal variance was used to determine no significance in total percentage of SCN7A+ surrounded neurons in underrepresented mice.

## Acknowledgements

Bioimaging Core Facility, Department of Biomedicine, Aarhus University

Imaging Core, Natural Science and Engineering Research Laboratory, University of Texas at Dallas. The authors thank the organ donors and their families for their gift of life.

---

**Supplementary Figure S1:**
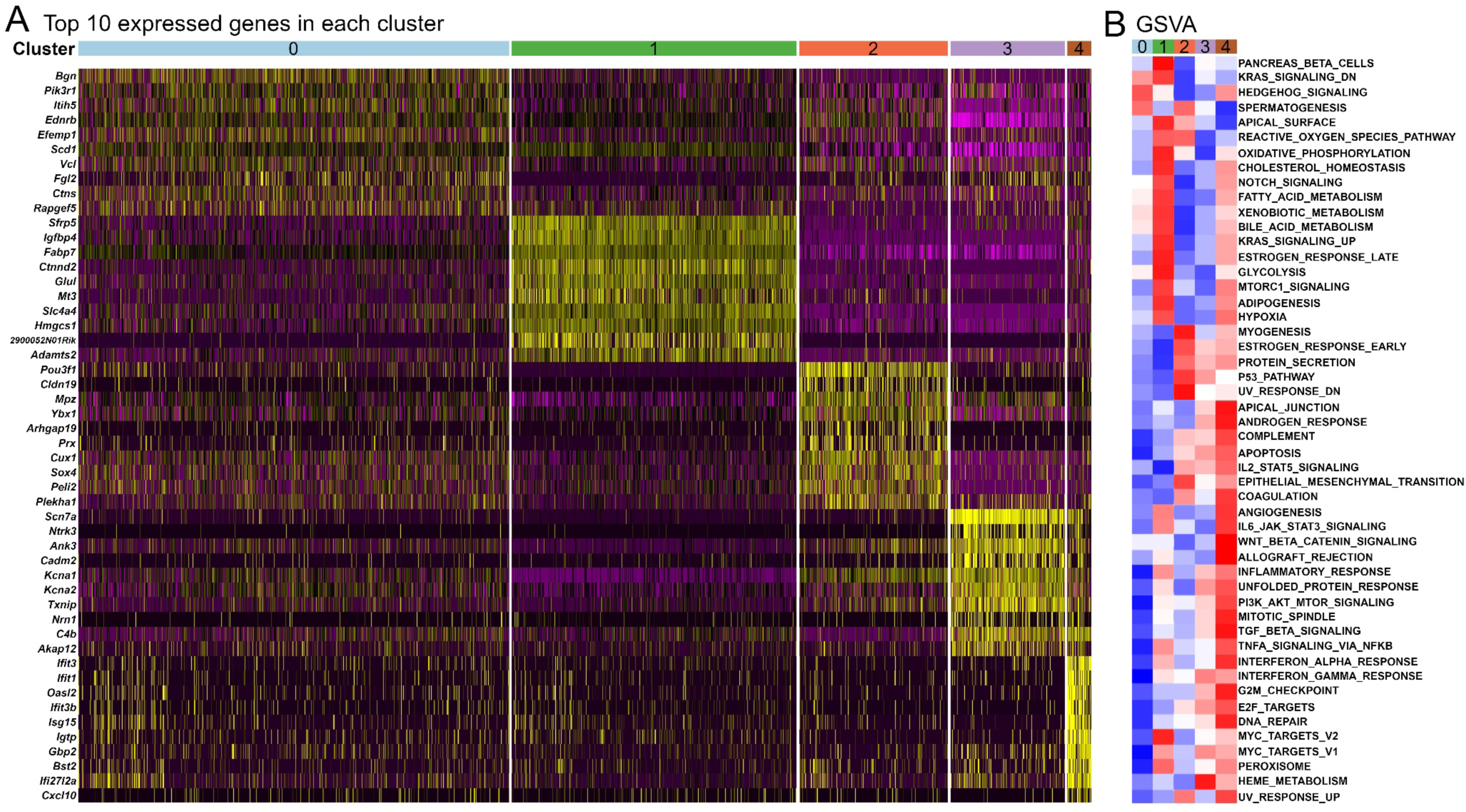
Genes expression and features of the identified subclusters. **A**: Top 10 expressed genes in each SGC subcluster. **B**: GSVA analysis of the SGC subclusters showing relative enrichment among biological functions.

**Supplementary figure S2:**
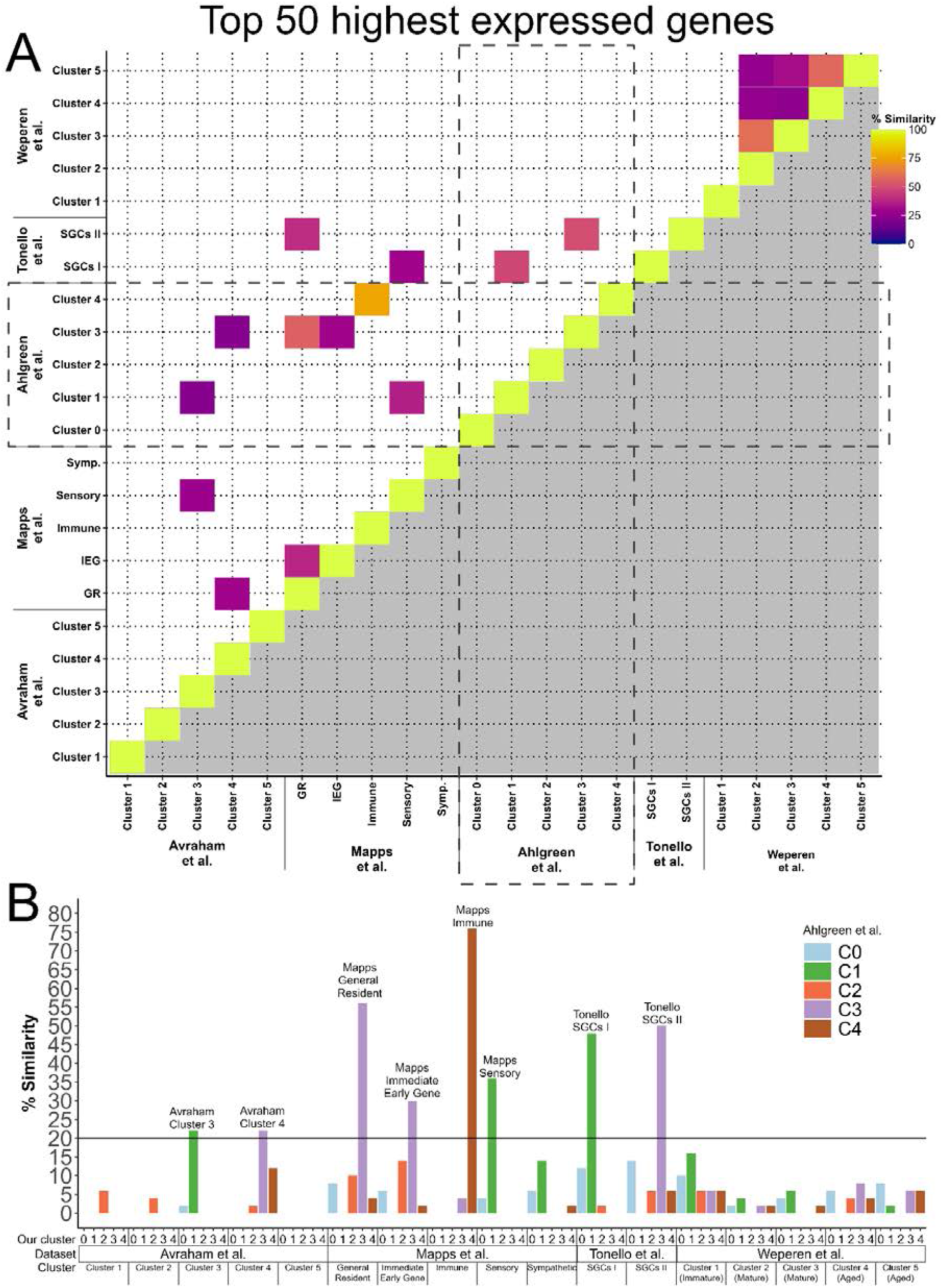
Comparison of the top 50 highest expressed genes among the clusters of five SGC heterogeneity studies. **A**: Comparison of similarity between the top 50 expressed genes of all clusters among published SGC heterogeneity studies with publicly available data and the present study. Threshold for inclusion: Avraham et al.: fold change > 1.5, p<0.05; Mapps et al.: fold change > 0.25, p.adj < 0.05; Tonello et al.: fold change > 1.25, p.adj<0.05; Weperen et al.: fold change > 0.25, p.adj<0.05; Ahlgreen et al.: fold change > 0.25, p.adj<0.05. Colored boxes indicate clusters with greater than 20% similarity. **B**: Bar plot showing percent similarity between top 50 expressed genes between clusters of Ahlgreen et al. and the already published SGC heterogeneity studies. Highlighted bars represent clusters among other datasets with over 20% similarity to a cluster in the present study and are found within the stippled line in panel (A).

**Supplementary figure S3:**
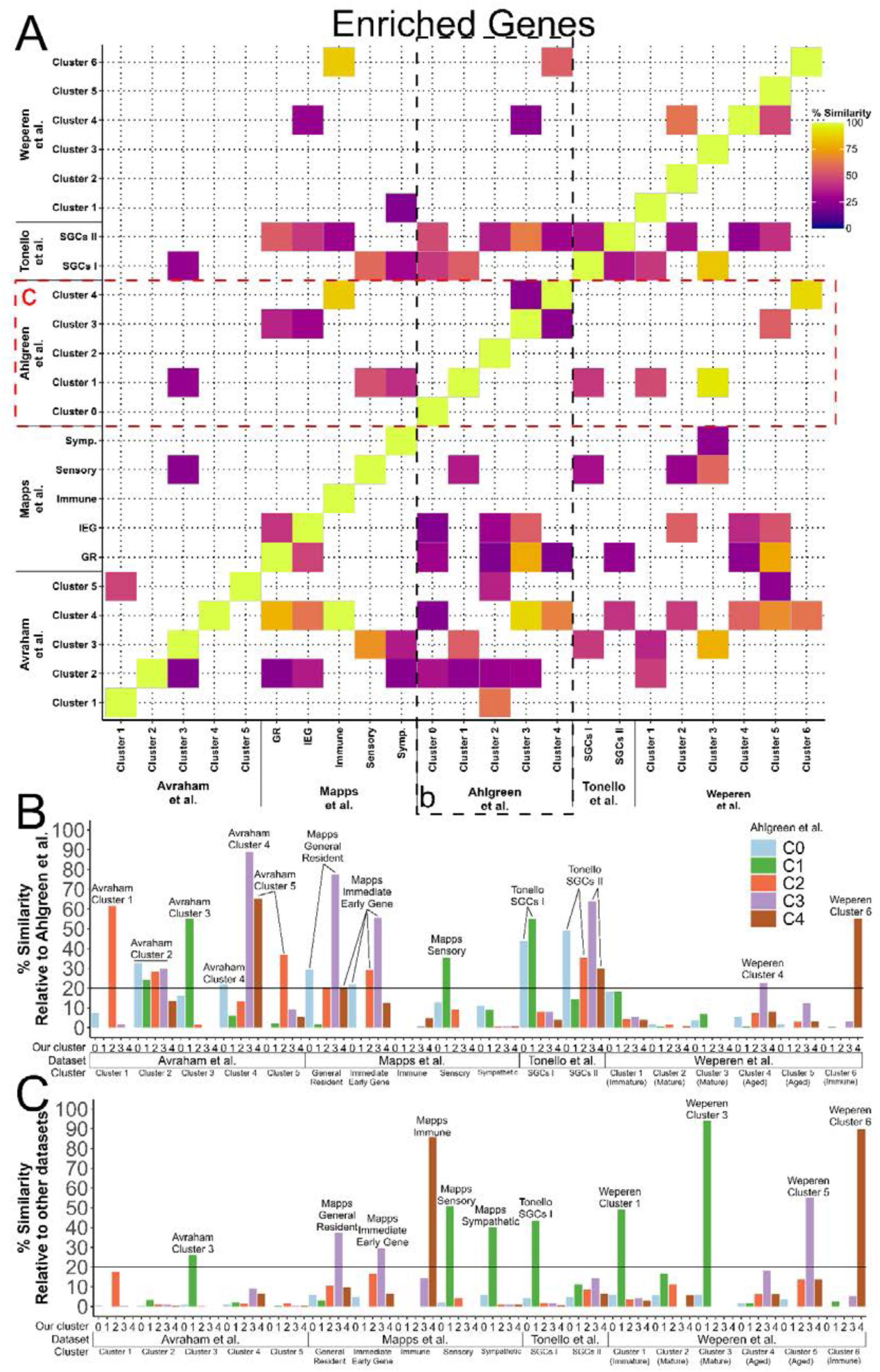
Comparison of enriched genes among the clusters of five SGC heterogeneity studies. **A**: Comparison of similarity between the enriched genes of all clusters among published SGC heterogeneity studies with publicly available data and the present study. Colored boxes indicate clusters with greater than 20% similarity. **B**: Bar plot showing percent similarity between enriched genes between clusters of Ahlgreen et al. and the already published SGC heterogeneity studies quantified relative to the number of enriched genes in the clusters of the other datasets. Highlighted bars represent clusters among other datasets with over 20% similarity to a cluster in the present study and are found within the black stippled line denoted (b) in panel (A). **C**: Bar plot showing percent similarity between enriched genes between clusters of Ahlgreen et al. and the already published SGC heterogeneity studies quantified relative to the number of enriched genes in the clusters of the present study. Highlighted bars represent clusters among other datasets with over 20% similarity to a cluster in the present study and are found within the black stippled line denoted (b) in panel (A).

**Supplementary figure S4:**
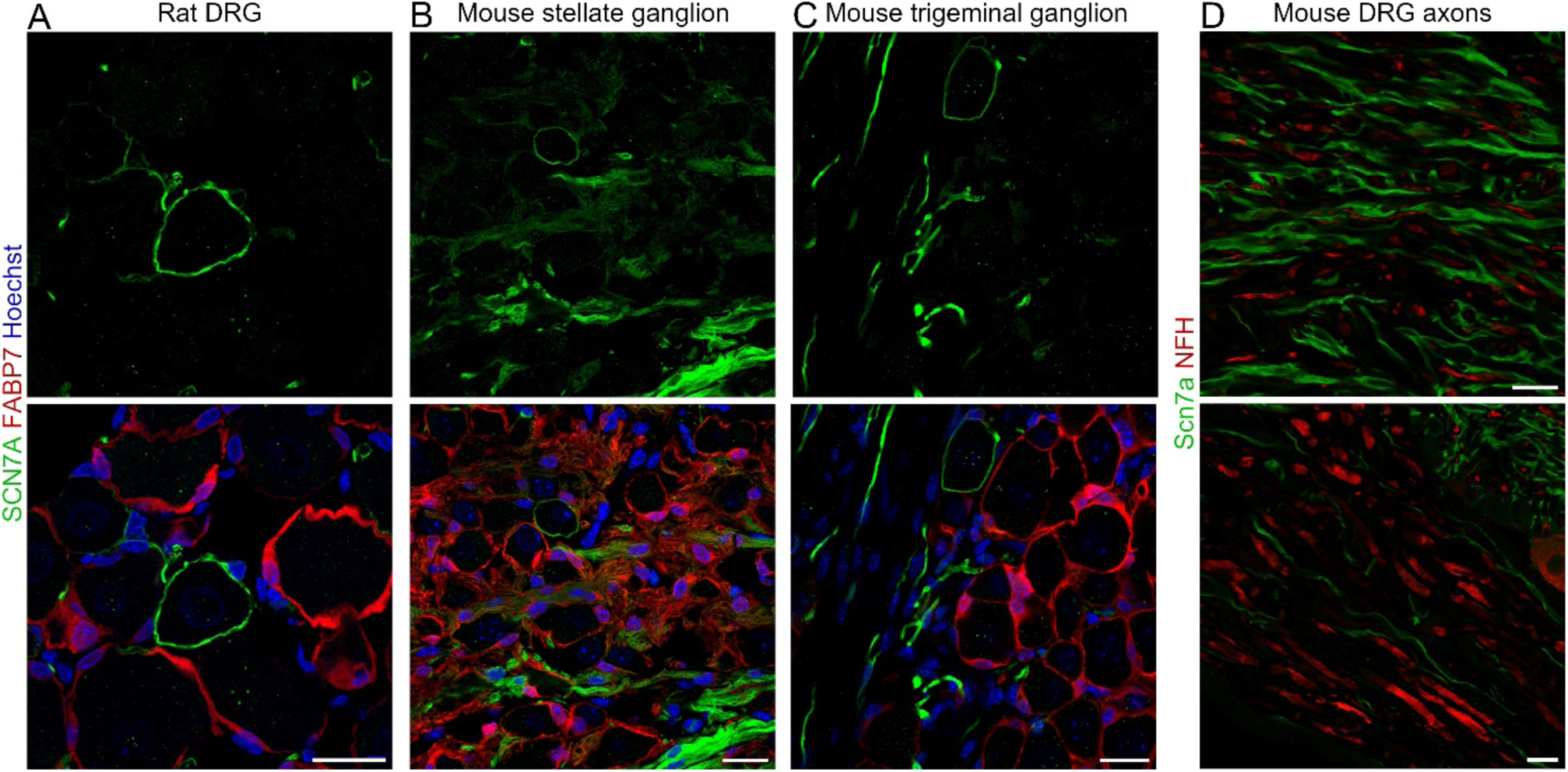
Further characterization of SCN7A+ SGCs. Immunohistochemical staining of different ganglia with SCN7A (green), FABP7 (red) and Hoechst (blue) showing the existence of SCN7A+ SGCs in **A**: rat DRG, **B**: mouse stellate (sympathetic) ganglion, and **C**: mouse trigeminal (sensory) ganglion. **D**: illustrates the disparate expression of SCN7A and NFH in mouse DRG axons. Scalebars: 20 µm.

**Supplementary Figure S5:**
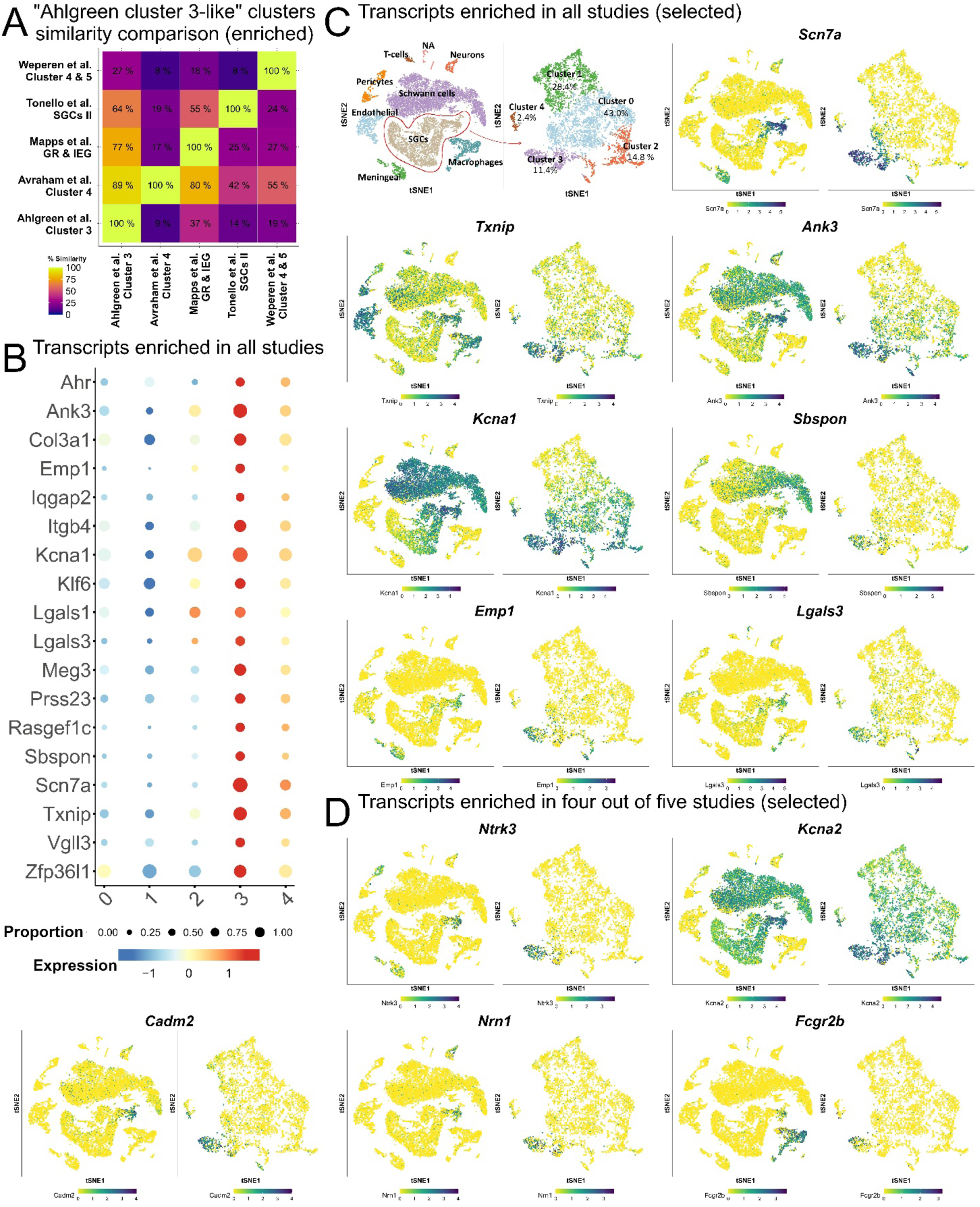
Comparative analysis of Scn7a-enriched clusters among datasets from published studies and the present study. **A**: Comparison of similarities between “Cluster 3-like” Clusters, defined as clusters enriched for Scn7a. In studies where multiple clusters displayed Scn7a-enrichment, these clusters were combined. From the Mapps et al. dataset, “General resident”- and “Immediate Early gene”-SGCs were combined (GR+IEG). From the Weperen et al. dataset, Cluster 4 and Cluster 5 were combined. Cluster 4 from the present study was also enriched for Scn7a, but was not combined with Cluster 3, given known differences in interferon response gene expression. **B**: Bubble plot displaying genes enriched in “Cluster 3-like”-clusters across all studies. 18 genes were common across studies. **C-D**: Selected genes enriched across either all studies (C) or 4 out of 5 studies (D) are displayed using t-SNE. Identification of an additional marker for SCN7A+ SGCs can be best narrowed down by studying the relative enrichment across SGC subtypes and other DRG cells.

**Supplementary Figure S6:**
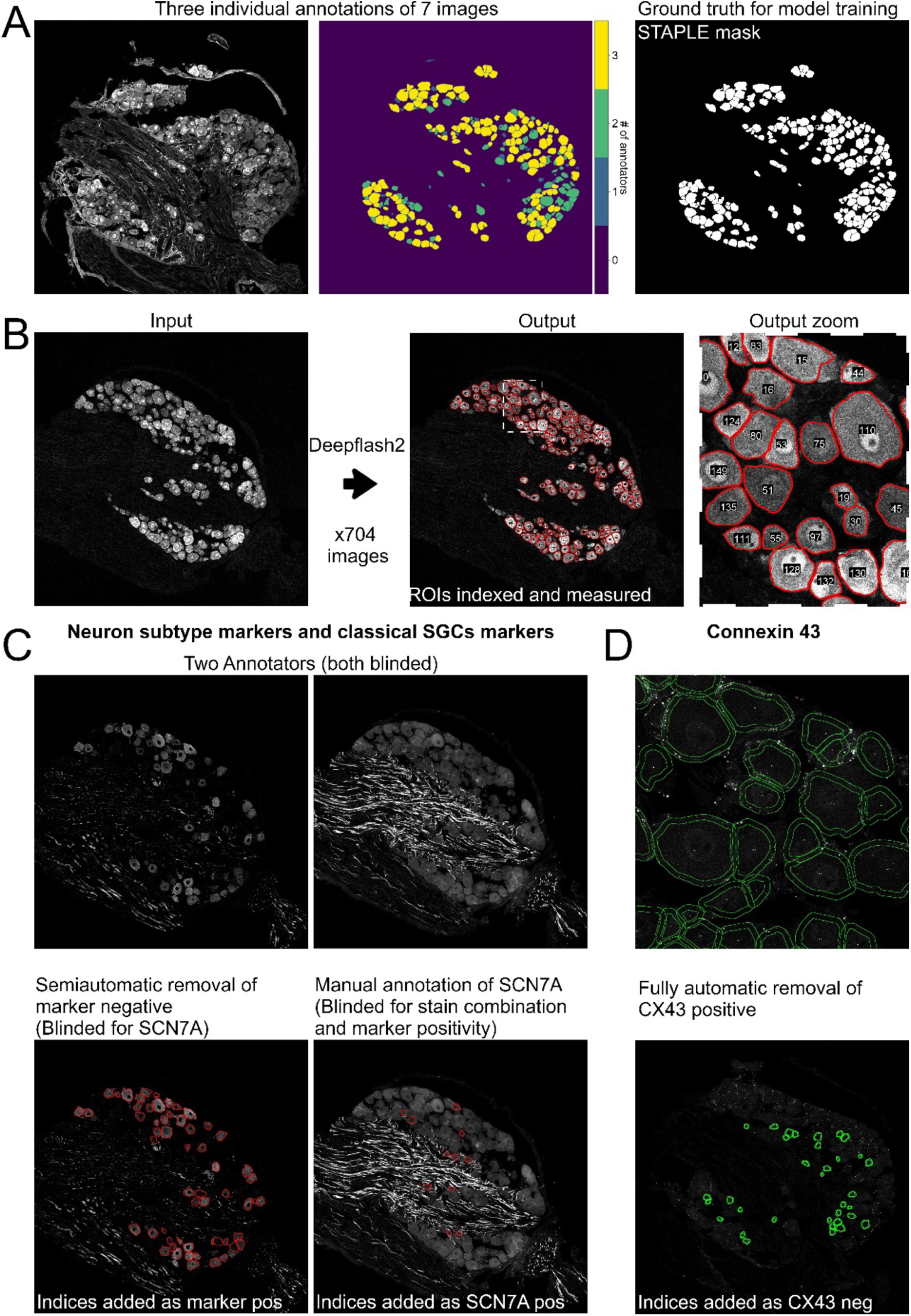
Automated annotation of neuronal somata in DRG sections using Deepflash2. **A**: Neuronal somata was manually annotated by three individuals on seven images. The STAPLE algorithm was used to determine the ground truth masks to be used for model training. **B**: Following the training of a five model deepflash2 ensemble, neuronal masks were predicted for the NEUN channel of 704 triple- or quadruple-stained DRG sections, and using Cellpose, instance segmented and converted to ROIs that were further processed in ImageJ. **C,D**: Output rois were applied to channels of SCN7A as well as neuronal and SGC markers and were labeled for positivity either semiautomatically using pixel values within the rois (neuronal and SGC markers), manually (SCN7A) or automatically using pixel values (CX43).

**Supplementary table S1:** Deepflash2 sample information. The combined dataset used for SCN7A+ SGC frequency quantification included a variable number of DRG sections representing each mouse, ranging between 4 and 42. No significant difference in percent SCN7A+ between under- and over-represented mice (unpaired 2-tailed students t-test with equal variance: p=0.83).

| **Sample** | **# Mice** | **#Unique**<br>**DRG/marker** | **#Unique**<br>**DRG/mouse** | **# Unique**<br>**neurons/marker** |
| --- | --- | --- | --- | --- |
| **Classical SGC markers** | **14** | **50.5±7.2** | **6.8±2.2** | **11570±1393** |
| FABP7 | 6 | 49 | 7-10 | 11676 |
| KIR4.1 | 6 | 55 | 7-11 | 11731 |
| CX43 | 6 | 57 | 8-11 | 13133 |
| GS | 8 | 41 | 4-7 | 9742 |
| **Neuron subtype**<br>**markers** | **14** | **49.3±4.6** | **6.4±2** | **10853±1865** |
| NFH | 6 | 50 | 7-9 | 10707 |
| CGRP | 6 | 46 | 5-9 | 9878 |
| SP | 6 | 52 | 8-9 | 11520 |
| TH | 6 | 51 | 7-9 | 12785 |
| IB4 | 6 | 52 | 8-9 | 11341 |
| Plexin C1 | 8 | 39 | 4-6 | 7300 |
| GFRα2 + IB4 | 8 | 43 | 4-7 | 9466 |
| OSMR + IB4 | 8 | 41 | 2-7 | 9640 |
| **Combined sample**<br>**unique neurons** | **#Mice** | **#Non-**<br>**overlapping**<br>**DRG sections** | **#Non-**<br>**overlapping DRG**<br>**sections/mouse** | **#Unique neurons** |
| **All neurons** | **14** | **214** | **4-42\*** | **48668** |

**Supplementary table S2:** Human donor demographics table. Abbreviations: CVA: cerebrovascular accident, MVA: motor vehicle accident.

| #Donor | Age | Sex | Ethnicity | Cause of Death |
| --- | --- | --- | --- | --- |
| 1 | 29 | Male | Black | Head trauma/blunt injury |
| 2 | 29 | Female | White | Anoxia/drug overdose |
| 3 | 27 | Male | Black | Anoxia/cardiac arrest |
| 4 | 31 | Female | White | CVA/stroke |
| 5 | 25 | Female | Black | Head trauma/blunt injury/MVA |
| 6 | 31 | Male | White | Anoxia/drug overdose |

**Supplementary table S3:**
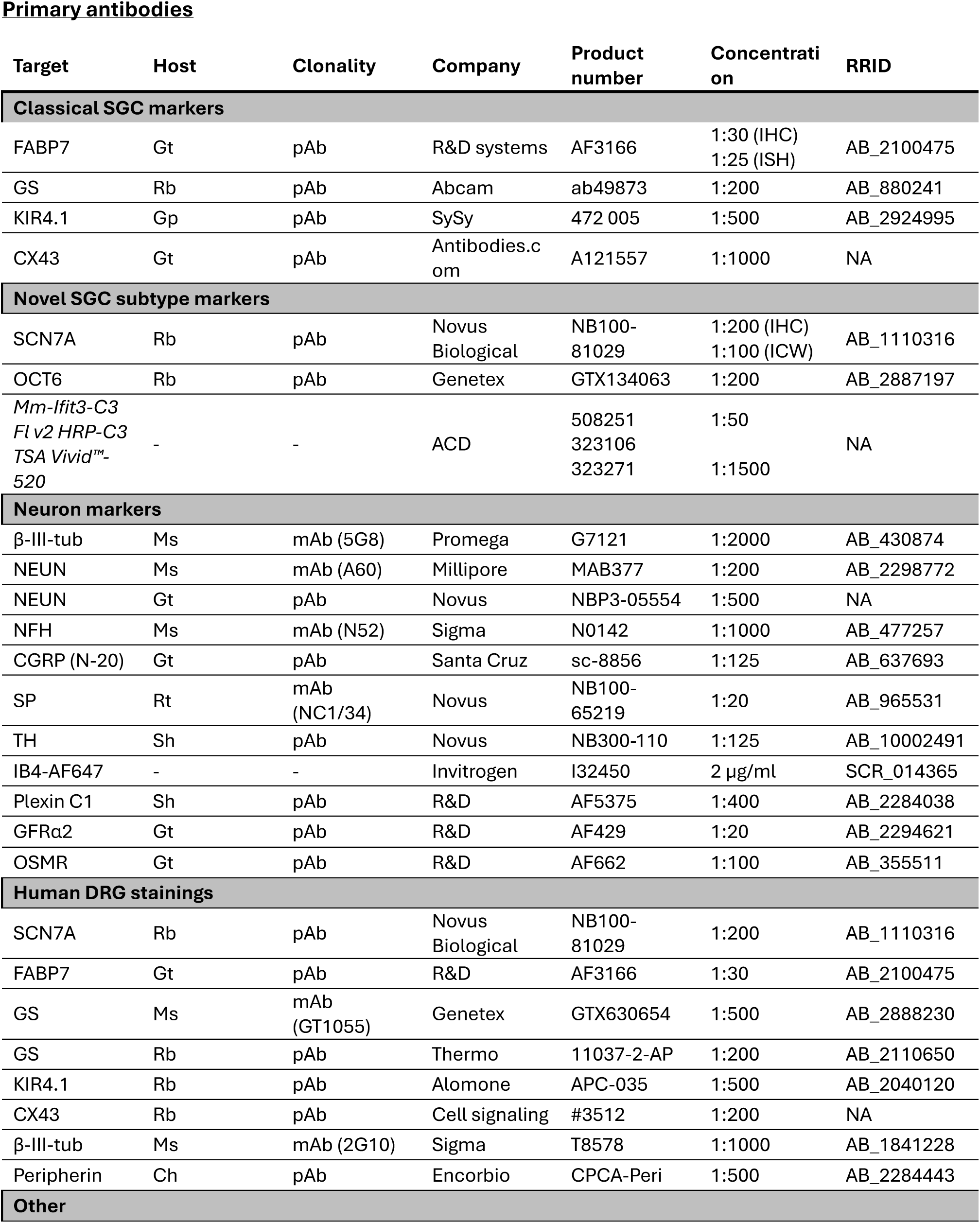
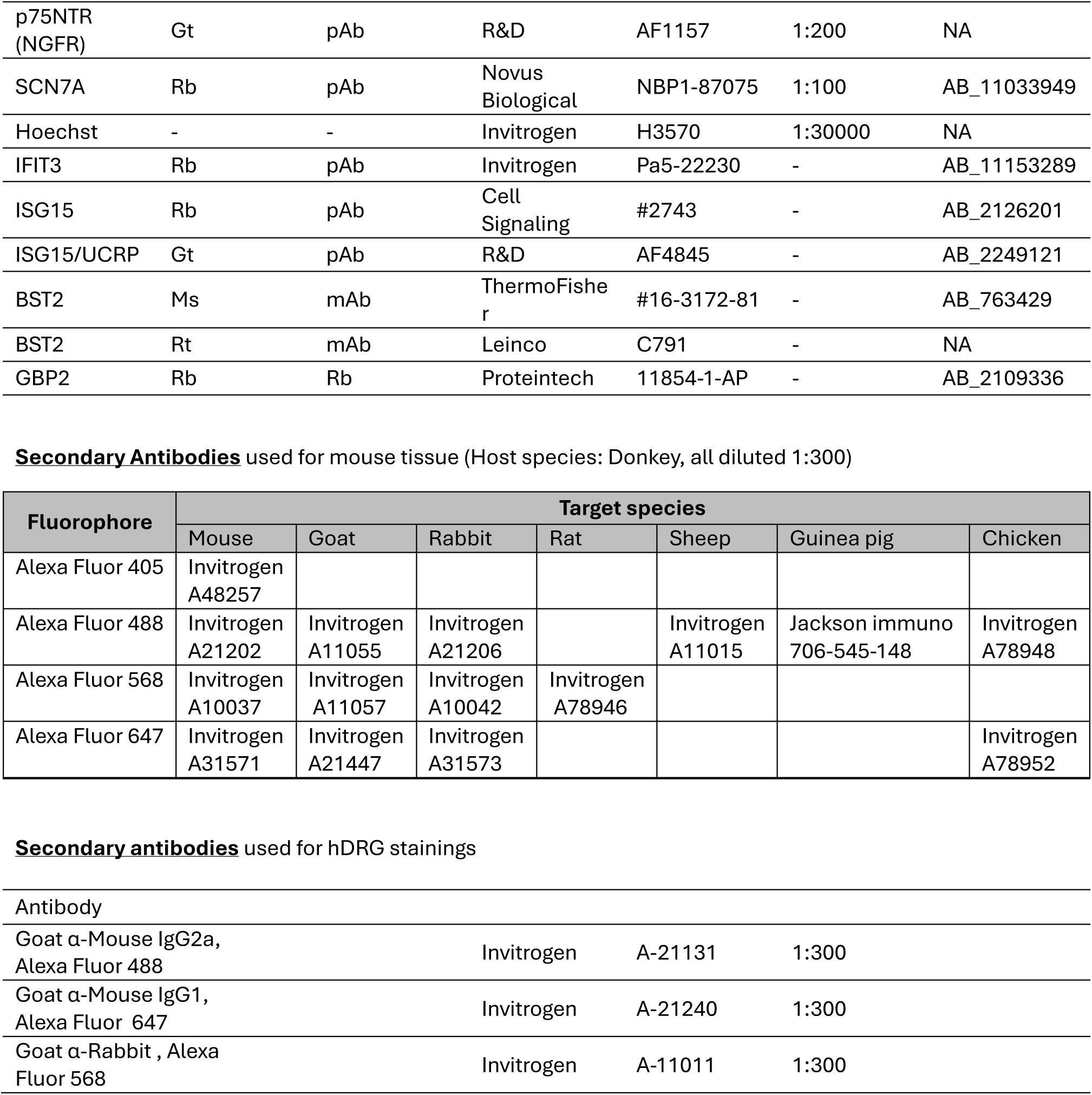
Antibodies and staining reagents. Abbreviations: * Gt: Goat – Ms: Mouse – Rb: Rabbit – Rt: Rat – Gp: Guinea pig – Sh: Sheep – Ch: Chicken – Dk: Donkey – pAb: polyclonal antibody – mAb: monoclonal antibody – IHC: immunohistochemistry – ICW: integrated co-detection workflow (in situ hybridization with IHC).

